# Reproductive fitness does not predict plastic changes in recombination due to diet in *Drosophila melanogaster*

**DOI:** 10.64898/2026.11.25.769255

**Authors:** Melika Ghasemi Shiran, Connor Haynes, Spencer Koury, Lauren McCann, Laney Hill, Natalia Rivera Rincón, Harlie Howard, Laurie Stevison

## Abstract

Meiotic recombination is a fundamental biological process that ensures proper chromosome segregation and generates novel genetic variation. Recombination rate exhibits plasticity in response to environmental conditions, yet how genetic background interacts with nutrition to influence this plasticity remains poorly understood. Using *Drosophila melanogaster* lines from the Drosophila Genetic Reference Panel (DGRP), we evaluated how varying caloric density affects X-chromosome recombination, body mass, fecundity, and survival across genotypes and over time. Caloric restriction significantly reduced female body mass in both genotypes, indicating a shared physiological response to dietary manipulation. Dietary treatment did not significantly affect egg-to-adult survival in marker-free crosses, whereas the high-calorie diet significantly reduced survival in marker-present crosses. After accounting for differential viability, we detected a significant genotype-by-environment (G×E) interaction in X-chromosome recombination rates. Furthermore, shifts in recombination rate were not associated with reduced survival or fecundity, failing to support adaptive hypotheses as explanations for recombination plasticity. To assess overall ovarian gene expression during these dietary perturbations, we conducted ovarian transcriptomic profiling (RNA-seq). Ovarian gene expression analysis revealed distinct, genotype-specific transcriptional responses to caloric restriction and excess calories, including significant changes in metabolic, stress-response, and cellular maintenance pathways across early and late time points. Overall, these findings demonstrate genotype-dependent recombination plasticity in response to dietary variation, without a consistent relationship between recombination rate and measured components of reproductive fitness, providing no support for the prediction of fitness-associated recombination under the dietary conditions examined. This work highlights the complex relationship of recombination rate plasticity among genotype, environment, and reproductive traits.

**Article Summary:** How does resource availability shape recombination plasticity, and does this response depend on genetic background? Using *Drosophila melanogaster*, we investigated how dietary caloric density influences meiotic recombination, reproductive fitness, and ovarian gene expression. We observed changes in body mass and substantial transcriptional responses to dietary variation, alongside genotype-specific recombination responses. After accounting for differential offspring viability, we found no consistent association between recombination rate and offspring survival. This contrasts with studies reporting associations between recombination plasticity and fitness. Our findings highlight the complexity of genotype-by-environment interactions and provide insights into the relationship between nutrition, reproductive fitness, and recombination plasticity.

## Introduction

Meiotic recombination is a core biological process that occurs during meiosis, involving the exchange of genetic material between homologous chromosomes (Petes 2001). This process creates novel combinations of alleles, contributing to genetic diversity within populations (Felsenstein 1974; Otto and Barton 1997; Otto and Lenormand 2002; Barton 2009). Reshuffling of existing genetic variation accelerates adaptation by reducing linkage disequilibrium, facilitating the accumulation of beneficial mutations, and promoting efficient selection across the genome (Otto and Barton 1997; Petes 2001; Otto and Lenormand 2002; Keightley and Otto 2006; Barton 2009). Additionally, recombination contributes to long-term genome evolution by affecting nucleotide diversity, genomic architecture, and the distribution of functional elements across chromosomes (Peñalba and Wolf 2020). Importantly, recombination is not a uniform process. Recombination rates exhibit substantial variation, varying among chromosomes, between sexes, and across environmental conditions (Petes 2001; Lenormand and Dutheil 2005; Modliszewski and Copenhaver 2017; Peñalba and Wolf 2020). Such variation suggests that recombination itself is an evolvable trait, shaped by both genetic and ecological factors, with significant consequences for evolutionary dynamics (Otto and Michalakis 1998; Lenormand 2003; Smukowski and Noor 2011; Stapley et al. 2017; Raffoux et al. 2026).

Beyond genetic variation, recombination rates display remarkable phenotypic plasticity with shifting external conditions. Individual recombination frequencies can change dynamically in response to physiological and environmental stressors encountered during maternal oogenesis (Stevison et al. 2017). Previous studies have demonstrated that recombination frequency can be influenced by factors such as temperature (Plough 1917), acute thermal stress (Jackson et al. 2015), pathogen exposure (Singh et al. 2015), and maternal age (Tedman-Aucoin and Agrawal 2012).

Furthermore, genetic backgrounds can differ in their average recombination rates (Hunter, Huang, et al. 2016). Genotype-by-environment interactions occur when the effect of an environmental condition on recombination differs between genetic backgrounds, producing different reaction norms. Nutritional status and energy availability influence reproductive physiology and germline development (Drummond-Barbosa and Spradling 2001). Although dietary effects on meiotic recombination remain relatively understudied, previous studies have shown that recombination rates can respond to dietary manipulation and that these responses can vary among genetic backgrounds in mice and *Drosophila melanogaster* (Belmonte-Tebar et al. 2022; Novak et al. 2026). *Drosophila melanogaster* is an ideal model for studying recombination plasticity due to its well-characterized genetics, short generation time, ease of dietary manipulation, and established recombination mapping tools (Ashburner and Bergman 2005; Comeron et al. 2012; Mackay et al. 2012). In Drosophila, recombination occurs primarily in females, being extremely rare in males. Recombination also exhibits substantial natural variation (Hunter, Huang, et al. 2016).

Caloric density refers to the amount of energy per unit of food and is a major determinant of physiological state and life-history traits in *Drosophila melanogaster*. Mechanistically, caloric intake may influence meiotic recombination via nutrient-sensing and metabolic signaling pathways that respond to energy status in *Drosophila melanogaster*. Nutrient availability modulates the insulin/insulin-like signaling (IIS) and Target of Rapamycin (TOR) pathways, which coordinate growth, energy balance, and reproductive physiology; these pathways also affect germline development and oogenesis in response to starvation or diet changes, thereby providing a physiological and signaling framework that could link caloric density to recombination outcomes (Hietakangas and Cohen 2009; Teleman 2009; Pritchett and McCall 2012; Smykal and Raikhel 2015; Nässel and Vanden Broeck 2016; Evangelakou et al. 2019). Diets with differing caloric densities alter traits such as body mass, reproductive outputs, and stress responses, and are commonly used to examine how energy availability influences organismal physiology and reproductive biology. For example, reducing caloric density has been shown to reduce fecundity while increasing lifespan and altering stress physiology in fruit flies (Partridge et al. 2005; Min et al. 2007). In the context of meiotic recombination, recent experimental manipulation of caloric density in *D. melanogaster* revealed that overall recombination rates increased as caloric intake decreased, with significant differences in crossover frequency between low-calorie (0.5×) and high-calorie (2×) diets in at least one genotype, indicating that caloric density can influence the crossover landscape along the X-chromosome (Novak et al. 2026). This effect was, however, strongly genotype dependent with temporal variation. Specifically, there were significant differences in crossover frequency between low-calorie (0.5×) and high-calorie (2×) diets in DGRP_42, but not in the DGRP_217 background, which did not show such plasticity. In DGRP_42, nutrient restriction led to increased recombination rate. Novak et al. (2026) also detected dietary effects on ovarian development, DNA fragmentation, and gene expression, with responses differing between genetic backgrounds. However, dietary treatment did not significantly affect fertility per backcross or the measured respiratory traits. These findings suggest that dietary effects on recombination should be interpreted alongside physiological measures rather than assuming a uniform decline in organismal performance.

Environmental effects on recombination raise questions about both the evolution of plasticity and its consequences for reproductive success. Models of fitness-associated and condition-dependent recombination examine circumstances under which selection can favor recombination rates that depend on an organism’s condition (Hadany and Beker 2003; Agrawal et al. 2005). However, observing an environmentally induced change in recombination does not by itself establish that the response evolved through selection or improves offspring fitness. Among the core hypotheses formalizing this adaptive view is the model of Fitness-Associated Recombination (FAR) (Hadany and Beker 2003; Agrawal et al. 2005). The fitness-associated recombination (FAR) hypothesis predicts a negative association between individual fitness and meiotic recombination rate. Under this theoretical framework, increased recombination in lower-fitness individuals may generate novel allelic combinations among offspring, potentially increasing the likelihood of producing higher-fitness genotypes (Hadany and Beker 2003; Agrawal et al. 2005). While this model is meant to explain the evolutionary advantage of sex, it also matters in the context of recombination rate plasticity. Empirical support for FAR in this context comes from studies showing increased recombination in stressed or low-fitness *Drosophila*. For example, exposure to acute heat stress in *D. melanogaster* is associated with both elevated recombination rates and reduced fitness, consistent with the negative correlation FAR predicts (Jackson et al. 2015). Similarly, desiccation-sensitive *Drosophila* genotypes subjected to dehydration stress show fitness-dependent increases in recombination and relaxation of crossover interference, with less fit genotypes producing more variable progeny relative to tolerant genotypes (Aggarwal et al. 2019).

Evidence for genotype-dependent recombination responses also extends beyond Drosophila. As summarized by Rybnikov et al. (2023), experiments in tomato found that temperature-induced changes in recombination depended on the genotype’s tolerance to temperature extremes, supporting a potential relationship between physiological sensitivity and recombination plasticity. Similarly, dietary manipulation in mice altered crossover-associated MLH1 foci in a strain-dependent manner (Belmonte-Tebar et al. 2022). Together, these studies suggest that the magnitude and direction of recombination responses depend on both genetic background and environmental perturbation. However, the small number of genetic backgrounds examined limits the generality of these findings and motivates broader sampling.

Previous work in *Drosophila melanogaster* demonstrated that DGRP_42 exhibited greater recombination rate plasticity and increased DNA damage during oogenesis with caloric restriction relative to DGRP_217 (Novak et al. 2026). These findings suggest a potential relationship between physiological stress and recombination plasticity. However, whether such responses are associated with reduced fitness, as predicted by FAR, remains unresolved. Theoretical advances further illustrate how fitness-dependent recombination can accelerate adaptation and maintain genetic diversity on rugged fitness landscapes (Hadany and Otto 2007) by favoring recombination in less fit individuals while preserving advantageous combinations in fit individuals.

Although maternal diet has been shown to modulate meiotic recombination rates, understanding whether these plastic responses represent adaptive, condition-dependent strategies remains challenging because directly linking recombination to fitness outcomes is methodologically difficult. Individual fitness in *Drosophila melanogaster* is conventionally quantified using downstream organismal traits such as maternal fecundity and egg-to-adult offspring survival (Clutton-Brock 1988; Fowler et al. 1997; Flatt 2020). However, these macro-level fitness proxies are frequently underpowered to isolate localized germline stress and are vulnerable to post-meiotic survival selection (Koury et al. 2026). In traditional testcross designs including recent evaluations of caloric density (Novak et al. 2026) recombination rates are typically inferred from visible marker counts without accounting for marker-specific viability effects, thereby confounding physiological stress responses with differential offspring survival. As a result, the relationship between nutritional stress, recombination plasticity, and individual fitness has not been directly measured, limiting our ability to interpret recombination changes as adaptive or condition-dependent responses.

Understanding how nutrition modulates recombination sheds light on how organisms regulate genetic variation in response to environmental conditions, with consequences for evolutionary potential and adaptive responses. More broadly, evaluating whether recombination plasticity represents an adaptive evolutionary strategy or a byproduct of physiological stress requires precise quantification of recombination rate variation across genotypes. Historically, multi-generation test crosses have been used to infer recombination rate plasticity. Conventional inference methods, however, rely primarily on scoring surviving F_2_ adult phenotypes rather than directly observing F_1_ meiocytes. This approach leads to substantial information loss and statistical confounding due to experimental mortality (Koury et al. 2026). Because only one of four chromatids per bivalent segregates to the functional egg pole during asymmetric female meiosis, and a substantial fraction of F_2_ individuals fail to reach adulthood under standard conditions, standard recombination metrics reflect only a small proportion of relevant meiotic events (Koury et al. 2026). This asymmetric loss introduces significant statistical uncertainty in estimates of recombination between treatments. Additionally, differential experimental mortality imposes pronounced selection bias on the observable dataset, especially when variation is driven by extreme genotypes or strong environmental perturbations such as nutritional stress (Singh et al. 2015; Hatkevich et al. 2017). Implementing a mechanistic framework that explicitly incorporates offspring survival and accounts for marker-specific survival effects allows for mathematical separation of germline recombination variation from downstream differential mortality (Koury et al. 2026). This approach enables higher statistical power and more accurate comparisons of reaction norms across environments and genotypes, reducing the historical risk of marker-induced survival distortions confounding meiotic estimates.

To resolve these historical limitations and build directly upon previous caloric density work in our lab (Novak et al. 2026), we conducted two pilot experiments to inform the design of the main experiment. In the first pilot experiment, we evaluated fecundity and egg-to-adult survival across six dietary caloric densities and selected three dietary treatments that provided sufficient offspring for reliable recombination rate estimation. In the second pilot experiment, we screened five genotypes across three selected dietary treatments to identify genotypes with contrasting recombination responses. Based on these preliminary results, we selected two genotypes with contrasting recombination responses across three dietary caloric densities to investigate genotype-by-environment interactions and evaluate whether variation in recombination rate is negatively associated with components of reproductive fitness, as predicted by the FAR hypothesis. We hypothesized that caloric restriction and caloric excess alter meiotic recombination rates and that the magnitude and direction of these responses differ among genotypes. Ultimately, resolving how nutritional variation interacts with genetic background to shape meiotic recombination provides critical insights into the mechanisms leading to genetic diversity and the adaptive potential of populations facing shifting environmental landscapes.

## Materials And Methods

### Fly Stocks and Husbandry

*Drosophila melanogaster* stocks were obtained from the Bloomington Drosophila Stock Center (BDSC). The genetic backgrounds used to assess the effect of genotype on recombination in the main experiment were DGRP_42 (BDSC 28127) and DGRP_399 (BDSC 25192) (Mackay et al. 2012). Also, genetic backgrounds used for the out crosses were *w1118* (BDSC 6326), Canton-S (BDSC 64349), and Oregon-R (BDSC 25211). To measure recombination, we used a multiply-marked X-chromosome carrying seven visible phenotypic markers: *yellow (y)*, *scute (sc)*, *cross-veinless (cv)*, *vermilion (v)*, *forked (f)*, *carnation (car)*, and *yellow+ (y^+^)*. Details regarding marker stock construction are provided in (Koury et al. 2026). The genomic identities and integrity of several DGRP genotypes were molecularly validated using a restriction fragment length polymorphism (RFLP) diagnostic protocol (Shiran et al. 2026).

Irrespective of dietary treatment, all stock maintenance, experimental crosses, and progeny rearing were conducted in a dedicated Percival incubator (Model DR41-VL) held strictly at 21°C, 70% relative humidity, under a controlled 12:12 hour light-dark cycle. To standardize parental and adult female densities, crosses used to generate experimental females were established with five females and five males per vial. Following eclosion, virgin females were maintained at a maximum density of 10 individuals per vial.

### Dietary Treatments and Food Manipulation

Experimental nutritional environments were created by altering the total caloric density of the standard laboratory diet (1.0×), which was based on the Bloomington Drosophila Stock Center formulation (Bloomington Drosophila Stock Center). Crosses were assigned to one of three dietary treatments: a low-calorie diet (0.5×), a standard control diet (1.0×), or a high-calorie diet (2.0×). To minimize technical variation, all food batches were prepared by the same investigator (LM).

This dietary design was based on the method of (Novak et al. 2026) with modifications to achieve precise scaling of caloric content while minimizing potential confounding effects of food consistency. For the 0.5× and 2.0× diets, the amounts of all calorie-containing ingredients (cornmeal, Brewer’s yeast, soy flour, and corn syrup) were reduced to half or increased to double those in the standard diet. This maintained the same relative proportions of macronutrients across all treatments.

All non-caloric components, including deionized water, propionic acid, Tegosept (methylparaben), and reagent alcohol, were kept constant to maintain preservative effectiveness and chemical stability. Agar was the only ingredient adjusted differently with careful consideration for maintaining similar food consistency across all diets and minimizing differences in texture. Its concentration was changed inversely with caloric density (6.2 g for 0.5×, 3.1 g for 1.0×, and 0.3 g for 2.0×). The complete basal diet formulations, including ingredient quantities and estimated caloric content (kcal per batch), are provided in Supplementary Table S1.

After the basal food solidified, 50 µL of yeast paste was applied to the surface of the food in each vial. Dry yeast quantities were scaled proportionally to the relative caloric density of the underlying diet. The composition and estimated caloric content of the supplemental yeast paste are provided in Supplementary Table S2. This supplementation differed from the experimental design of Novak et al. (2026), in which no supplemental yeast paste was provided.

### Genetic Crosses and Experimental Design

To map crossovers and infer recombination frequencies across nearly the entirety of the X-chromosome, we employed a modified version of the six-point testcross design detailed in (Koury et al. 2026). This study focuses on two *Drosophila melanogaster* genotypes exposed to three dietary treatments. The design of this experiment was informed by two pilot experiments (see Supplementary Methods for detailed descriptions). First, a pilot using a single genotype across 6 caloric densities was used to evaluate the physiological limits, fecundity, and egg-to-adult survival across six dietary treatments. This first pilot showed that the extreme diets used in (Novak et al. 2026) did not significantly reduce egg-to-adult survival as compared to the intermediate caloric densities. A second pilot experiment (5 genotypes × 3 environments, two replicates each) screened five wild-type DGRP genotypes across the three diets selected from the first pilot and identified two genotypes with contrasting recombination responses to diet. These two genotypes were selected for the fully powered experiment described here.

Parental (P) crosses were established by placing five mutant females with five wild-type males on one of the three assigned dietary treatments. Between two and nine parental replicate vials were established for each genotype and diet combination. Parents remained on their assigned diets throughout mating and egg laying so that F_1_ offspring experienced continuous exposure to their assigned diet from the egg stage through larval and pupal development until adult eclosion (emergence from the pupa).

Virgin F_1_ females were collected within 8 hours of eclosion and transferred to fresh vials containing the same dietary treatment, where they were maintained for 1–8 days to reach sexual maturity. To verify virginity, holding vials were monitored for three weeks, and any vial containing larvae, pupae, or other evidence of unintended mating was discarded.

Sexually mature F_1_ virgin females, carrying either the multiply marked X-chromosome or the marker-free X-chromosome, were backcrossed to males of the corresponding genotype that had experienced the same dietary treatment. Each backcross consisted of three females and three males per vial. The detailed genetic crossing scheme is illustrated in Figure 2.

**Figure 1.**
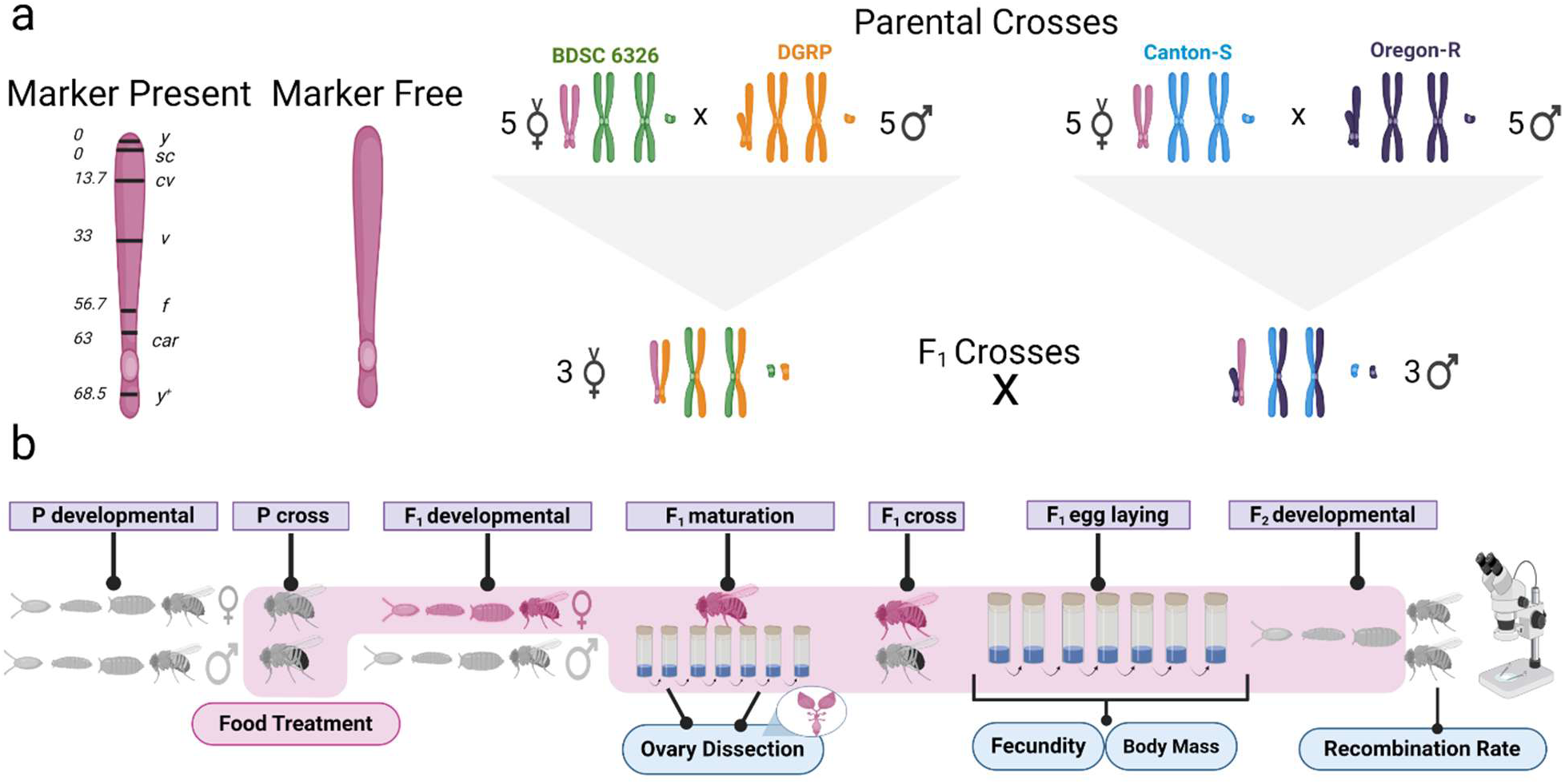
Experimental Design and Crossing Schemes. Two types of genetic crosses were conducted – marker present and marker free (a) genetic backgrounds across Drosophila melanogaster strains (BDSC 6326, DGRP, Canton-S, and Oregon-R), followed by F1 crosses. The marker-present X-chromosome indicates the physical locations of visible markers (y, sc, cv, v, f, car, y⁺). (b) Timeline of developmental and experimental phases, detailing timing of dietary treatments , ovary dissection, body mass assays, fecundity measurements across daily transferred broods, and F2 recombination rate estimation. Created in BioRender. Shiran, M. (2027) https://BioRender.com/xpmici9

**Figure 2.**
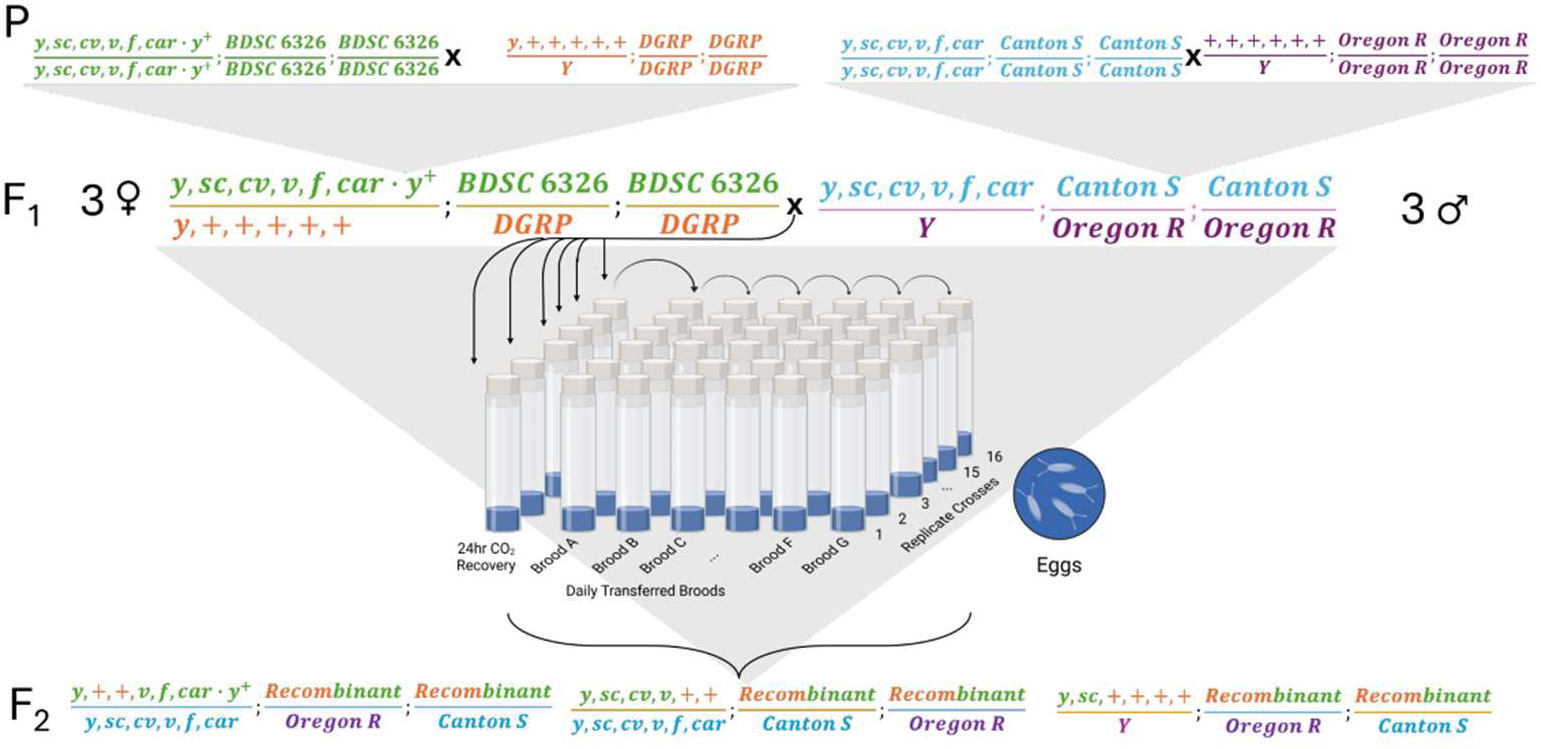
Detailed genetic crossing scheme for F1 and F2 offspring generation. Crossing diagram illustrating parental (P), F1, and F2 generation genotypes across daily transferred broods (Broods A–G) following 24-hour CO₂ recovery. Shows the inheritance of marker-free and marker-present chromosomes and resulting F2 recombinant genotypes paired with Canton-S, Oregon-R, and Y-chromosome components. Created in BioRender. Shiran, M. (2026) https://BioRender.com/i06w0fk.

Sixteen independent biological replicates were established for each genotype × diet combination in the marker-present crosses and four biological replicates for the marker-free controls. After a standardized 24-hour recovery period following CO₂ anesthesia (Barron 2000; Nilson et al. 2006), mated females were transferred to fresh treatment-specific food. They were subsequently tapped transferred every 24 hours for seven consecutive days, generating seven sequential broods. This design produced 672 marker-present F_2_ progeny vials (2 genotypes × 3 diets × 16 replicates × 7 broods) and 168 marker-free F_2_ progeny vials (2 genotypes × 3 diets × 4 replicates × 7 broods), for a total of 840 F_2_ progeny vials used for recombination analysis.

### Body Mass Measurement

Body mass was measured exclusively in the F_1_ females over the seven-day experimental window. To maintain handling parity with the other fitness assays, F_1_ females were transferred to fresh food every 24 hours. From each vial, three females were randomly selected for weighing and each female weighed separately. Flies were anesthetized using the FlyNap® Anesthetic Kit and weighed using a Sartorius Model 1712 MP8 Silver Edition analytical balance. Following weighing, these females were discarded and were not returned to the experiment or used in any subsequent assays.

### Ovary Dissection

To capture the molecular transcriptomic impacts of caloric density during oogenesis, F_1_ ovaries were harvested at two specific time points: 2 days and 6 days post-eclosion. Samples were collected from marker-free, virgin F_1_ experimental females for both genotypes (DGRP_42 and DGRP_399) across all three dietary environments (0.5x, 1x, and 2x).

All dissection tools, stereomicroscopes, and lab work surfaces were thoroughly treated with an RNase decontamination solution (RNase Away) to maintain strict RNase-free conditions. Flies were anesthetized via CO_2_ for no longer than ten minutes during processing to minimize stress-induced transcriptional artifacts. Ovaries attached to the oviduct were cleanly removed by gently stabilizing the fly thorax with ultra-fine forceps while pulling the posterior abdomen under a stereo microscope. For each distinct condition, a biological sample pooling ten pairs of ovaries (from 10 individual females) was generated. Each pool was immediately transferred into pre-chilled tubes containing 100 µl of DNA/RNA Shield (Zymo Research), held briefly on ice during the processing window, and stored at -80°C until subsequent extraction. Two independent biological replicates were prepared per genotype, diet, and timepoint combination (N_re*p*_ = 2), generating a total of 24 sequenced ovarian libraries.

### RNA Extraction and Quality Assessment

Total RNA extraction was executed utilizing the Direct-zol RNA Miniprep Plus Kit (Zymo Research, Kit #R2071-A) following the manufacturer’s protocol. An on-column DNase I treatment was performed during RNA extraction to remove contaminating genomic DNA. Extracted total RNA was initially evaluated for raw concentration and purity via an indexing spectrophotometer (Thermo Fisher Scientific NanoDrop), requiring absorbance ratios (260/280 and 260/230) strictly greater than 2.0 for sample progression.

To satisfy downstream high-throughput requirements, RNA pools were further purified and concentrated utilizing the RNA Clean & Concentrator-25 Kit (Zymo Research, Kit #R1017). Final concentrations were re-verified via NanoDrop and confirmed using a Qubit RNA BR Assay Kit (Thermo Fisher Scientific). Independent quality control validation was conducted by Novogene via agarose gel electrophoresis and capillary electrophoresis profiling on an Agilent 2100 Bioanalyzer. Only high-integrity samples exhibiting an RNA Integrity Number (RIN > 9.0) were advanced to high-throughput library preparation.

### RNA-seq Library Preparation and Sequencing

High-throughput transcriptomic library construction, raw sequencing services, and differential expression analysis were executed by Novogene. Strand-specific, poly-A selected mRNA libraries were prepared from qualified total RNA inputs using standard Illumina-compatible workflows. The resulting 24 multiplexed libraries were sequenced on an Illumina platform using 150bp paired-end (PE150) chemistry, targeting an average sequencing depth of approximately 42–82 million raw reads per individual library sample.

Raw fastq datasets were processed through a standardized bioinformatic pipeline to check sequencing quality. Clean reads were isolated by filtering out low-quality reads, adapter sequences, and heavily poly-N-contaminated fragments, maintaining high-quality thresholds (targeting sequencing error rates ∼ 0.01%, and high Q20 and Q30base metrics). Filtered, clean paired-end reads were uniquely mapped against the *Drosophila melanogaster* reference genome (BDGP6, GenBank assembly accession GCA_000001215_4) utilizing HISAT2 (v2.2.1). Raw gene-level fragments and transcript abundances were quantified programmatically using feature Counts (v2.1.1). High consistency among biological replicates was confirmed by squared Pearson correlation metrics (r^2^ = 0.980–0.992) prior to differential gene expression analysis.

### RNA-seq Statistical Analysis

Differential gene expression analysis was performed by Novogene using the ‘DESeq2’ package (v1.42.0; Love et al. 2014), which fits negative binomial generalized linear models to gene-level read counts. Statistical significance was assessed using Wald tests, and *p*-values were adjusted for multiple testing using the Benjamini–Hochberg procedure. Pairwise dietary comparisons included high-calorie (2×) versus control (1×), low-calorie (0.5×) versus control (1×), and low-calorie (0.5×) versus high-calorie (2×) treatments, both across pooled samples and separately within each combination of genetic background and sampling day. Differential expression results provided by Novogene were imported into R for downstream analysis. Genes with missing adjusted *p*-values or log₂ fold changes were excluded, and genes were classified as differentially expressed (DEGs) using an FDR-adjusted *p*-*value* < 0.05 and an absolute log₂ fold change > 1.

Downstream biological systems testing and pathway mapping were executed via the ‘clusterProfiler’ (v4.8.1) package. Statistical enrichment tracking for Gene Ontology (GO) terms (including Biological Process, Molecular Function, and Cellular Component) and Kyoto Encyclopedia of Genes and Genomes (KEGG) molecular pathways was determined utilizing a significance cutoff of padj ≤ 0.05 after Benjamini-Hochberg correction. Transcriptomic intersection patterns, expression profiles, and pathway signals were visualized in R using the ‘UpSetR’ (Conway et al. 2017), ‘ggplot2’ (Wickham 2016), and ‘ggnewscale’ (Campitelli 2026) software packages.

### Fecundity Assay

Daily F_1_ female fecundity was quantified over the 7-day period using an absolute egg-laying assay adapted from (Schneider et al. 2025). Experimental vials were photographed within 24 hours of fly transfer prior to hatching. Egg counts were performed directly from these digital images by two independent researchers (CH and LM), as well as a third investigator (LH) who analyzed pilot data. Eggs deposited in obscure regions not fully visible in the photographs (e.g., attached to vial walls, embedded in food crevices, or inside small holes) were quantified manually via visual inspection under a microscope.

Immediately following this imaging and manual wall-egg counting process, a 15 cm piece of Kimwipe was inserted into the media. Paper was wetted with 100 µL of an acid solution containing 0.01M propionic acid and 0.001M phosphoric acid to avoid microbial and mold growth during larval development.

To correct for systematic image-based undercounting, a predictive ordinary least squares (OLS) linear regression model was constructed independently for each of the three investigators mapping raw digital image-based counts to corresponding manually validated egg counts obtained from a separate calibration dataset not associated with the focal experimental treatments (Schneider et al. 2025). These distinct regression equations were applied to systematically adjust the raw counts for each observer across the entire experimental dataset (Supplementary Figure S1). Following calibration, the two adjusted egg counts were averaged for each vial and divided by the number of females to estimate daily fecundity (eggs per female per 24 h).

Daily F₁ female fecundity was analyzed separately for marker-free and marker-present crosses using linear mixed-effects models (Fecundity ∼ Treatment * Genotype * Brood + (1 | vialID)). Treatment, genotype, brood, and all their interactions were included as fixed effects, and vial identity was included as a random intercept to account for repeated measurements across broods. Because fecundity was quantified before adult offspring emerged and was based on the mean of two adjusted egg counts, adult offspring observer identity was not included in the fecundity models. Statistical significance was assessed using Type III F-tests with Satterthwaite’s approximation for denominator degrees of freedom. Pairwise dietary comparisons within each genotype were performed using estimated marginal means with Tukey adjustment for multiple comparisons.

### Survival Calculation

Adult survival was monitored daily following the initiation of adult eclosion. Adults were anesthetized using CO_2_ and separated based on sex and/or phenotype under a Nikon SMZ645 Stereo Microscope or an Olympus SZ61 microscope. Newly eclosed adults were counted daily for approximately two weeks. Specifically, counting continued until no new adults emerged for three consecutive days, ensuring complete emergence of the cohort and preventing generational overlap. Vials from each genotype and treatment combination were randomly distributed during counting of adults to minimize observer bias. Survival was calculated as the proportion of initial eggs (including both surface and wall eggs tracked during the fecundity assay prior to Kimwiping) that successfully developed into adults.

Egg-to-adult survival was analyzed separately for marker-free and marker-present crosses using linear mixed-effects models (Survival ∼ Treatment * Genotype * Brood + (1 | vialID) + (1 | Data_collector)). Treatment, genotype, brood, and all their interactions were included as fixed effects, with vial identity and adult offspring observer identity included as random intercepts to account for repeated measurements and potential differences in offspring scoring, respectively. Statistical significance was assessed using Type III F-tests with Satterthwaite’s approximation for denominator degrees of freedom.

### Recombination Mapping and Multi-Locus Estimation Framework

Recombination frequencies and meiotic crossover dynamics across the X-chromosome were quantified from the transmission of visible genetic markers to the progeny following established methodology (Koury et al. 2026, see below). Emerging offspring from the multiply marked crosses were collected daily, anesthetized via CO_2_, and phenotypically classified under a Nikon SMZ645 or Olympus SZ61 stereo microscope for approximately two weeks post-eclosion to capture the complete cohort. Recombination was estimated using two complementary approaches: observed recombination fractions converted to Kosambi map distances and an extended counting model incorporating differential progeny viability.

#### 1. Empirical Recombination Frequency Calculation

First, empirical interval-specific and whole-chromosome recombination frequencies (*r*) were calculated directly from raw phenotypic counts. Offspring were classified into recombinant or non-recombinant classes based on the presence or absence of the seven visible X-linked genetic markers (*y*, *sc*, *cv*, *v*, *f*, *car*, and *y+*). The empirical recombination frequency was mathematically defined as the total number of observed recombinant events within a given interval divided by the total number of scored progeny. Raw recombination fractions were subsequently converted into genetic map distances in centiMorgans (cM) using Kosambi’s map function (Kosambi 1943) to correct for crossover interference across individual intervals. This approach characterizes observed recombination across the assayed intervals without explicitly accounting for differential survival among progeny classes.

#### 2. Multi-Locus Data-Generating Process and Counting Model Fitting

Second, to account for systematic mutant survival effects, sex-specific adult phenotypic class counts from each experimental unit were fit to an extended counting model of crossover patterning as detailed in Koury et al 2026. This framework models recombination as a multi-step data-generating process assuming that: (1) Programmed double-strand breaks occur as a homogeneous Poisson point process along the genetic map; (2) crossover maturation is a stationary renewal process; (3) chromosome segregation represents a random one-half thinning of this process; (4) fertilization is Mendelian; and (5) egg-to-adult survival is binomially distributed with a sex-and treatment-specific rate parameter determined by the unique complement of marker alleles.

Maximum likelihood estimates of the model parameters, including double-strand break density, genetic lengths (L, in centiMorgans), and crossover interference strength (m), were obtained using a downhill simplex optimization method implemented in custom scripts in R version 4.5.3 (R Core Team 2026), using the ‘dfoptim’ package version 2023.1.0 (Varadhan et al. 2023). Survival parameters were constrained between 0 and 1.

Prior to statistical analysis, outliers in both Kosambi and viability-corrected genetic map length estimates were identified and excluded using the 1.5 × interquartile range (IQR) criterion. For each estimation method, observations below Q1 − 1.5 × IQR or above Q3 + 1.5 × IQR were excluded, with thresholds calculated across all genotypes, dietary treatments, and broods. This procedure resulted in the exclusion of 15 of 653 (2.30%) non-missing Kosambi estimates and 18 of 659 (2.73%) viability-corrected estimates. Additionally, six observations with missing Kosambi estimates were excluded, resulting in 638 and 641 observations retained for statistical analysis, respectively.

### Estimation of Crossover Interference

Crossover interference was estimated using the Cₓ(Cₒ)ᵐ counting model (Zhao et al. 1995; Koury et al. 2026), where *m* represents the number of noncrossover events between successive crossover events. Maximum likelihood estimates of the renewal-process parameter (*p* = *m* + 1) were obtained from pooled marker-present cross data. Data were pooled by adult offspring observer, dietary treatment (0.5×, 1.0×, or 2.0×), and genotype (DGRP_42 and DGRP_399), producing four replicate estimates per treatment × genotype combination (*n* = 24). Crossover interference estimates were analyzed using a two-way ANOVA (MLE_p ∼ Diet * Genotype).

## Results

Here, we compared X-chromosome meiotic recombination under varying caloric densities across genotypes, with a focus on the relationship with fitness. We used visible markers to measure recombination rate in a traditional 3-generation genetic cross. In total, the 288 F₁ females laid a total of 46,798 eggs, yielding 29,562 F₂ progeny that were scored to infer recombination (Table 1). Additionally, we conducted a marker-free cross to compare fitness differences due to the treatments to fitness differences due to the visible markers. In total, there were 72 F_1_ females which laid 22,461 eggs, yielding an additional 14,762 F_2_ progeny. The marker free crosses were also used to measure the impact of the treatments in the absence of visible markers on both body mass and ovarian transcription.

**Table 1.** Summary of reproductive fitness, offspring survival, map length, and crossover interference across genotypes and dietary treatments. Total F₁ female fecundity, mean F₂ egg-to-adult survival, X-chromosome genetic map length (cM), and crossover interference are presented for two Drosophila melanogaster genetic lines (DGRP_42 and DGRP_399) maintained under three dietary caloric densities (0.5×, 1.0×, and 2.0×) in marker-free and marker-present crosses. Crossover interference is reported as the mean estimated renewal-process parameter (p = m + 1) of the Cₓ(Cₒ)ᵐ counting model, where m represents the number of noncrossover events between successive crossover events.

|  |  | Marker free cross |  | Multiply marked cross |  |  |  |
|---|---|---|---|---|---|---|---|
|  |  | Total F<sub>1</sub><br>Fecundity | Mean F<sub>2</sub><br>Survival | Total F<sub>1</sub><br>Fecundity | Mean F<sub>2</sub><br>Survival | X Chr.<br>Length | Interference |
| 42 | 0.5x | 3125 | 72% | 4401 | 73% | 65.9 | 3.25 |
|  | 1.0x | 3701 | 72% | 7015 | 78% | 67.6 | 3.25 |
|  | 2.0x | 4000 | 82% | 5949 | 57% | 69.2 | 3.5 |
| 399 | 0.5x | 3919 | 45% | 8324 | 74% | 78.6 | 3.25 |
|  | 1.0x | 3692 | 69% | 9384 | 65% | 72.1 | 2.75 |
|  | 2.0x | 4024 | 58% | 11725 | 49% | 73.3 | 4.75 |

### Caloric Restriction Reduces Female Body Mass in Both Genotypes

Mean adult female body increased with dietary caloric density in both genotypes, with DGRP_42 consistently exhibiting greater mean body mass than DGRP_399 across all dietary treatments (**Figure 3**a). A linear mixed-effects model revealed a significant effect of dietary treatment on female body mass (*F*₂,₁₁₉.₈₅ = 34.46, *p*-value = 1.50 × 10⁻¹²). Genotype also significantly affected body mass (*F*₁,₁₂₀.₀₅ = 9.17, *p*-value = 0.003), with DGRP_42 exhibiting greater mean body mass than DGRP_399. Body mass also varied significantly among broods (*F*₆,₁₁₉.₂₅ = 5.72, *p*-value = 2.94 × 10⁻⁵). However, neither the treatment-by-genotype interaction (*F*₂,₁₁₉.₈₅ = 0.70, *p*-value = 0.501), the treatment-by-brood interaction (*F*₁₂,₁₁₈.₉₄ = 1.69, *p*-value = 0.076), the genotype-by-brood interaction (*F*₆,₁₁₉.₂₅ = 1.76, *p*-value = 0.114), nor the three-way interaction (*F*₁₂,₁₁₈.₉₄ = 0.99, *p*-value = 0.462) was significant (Table 2).

**Figure 3.**
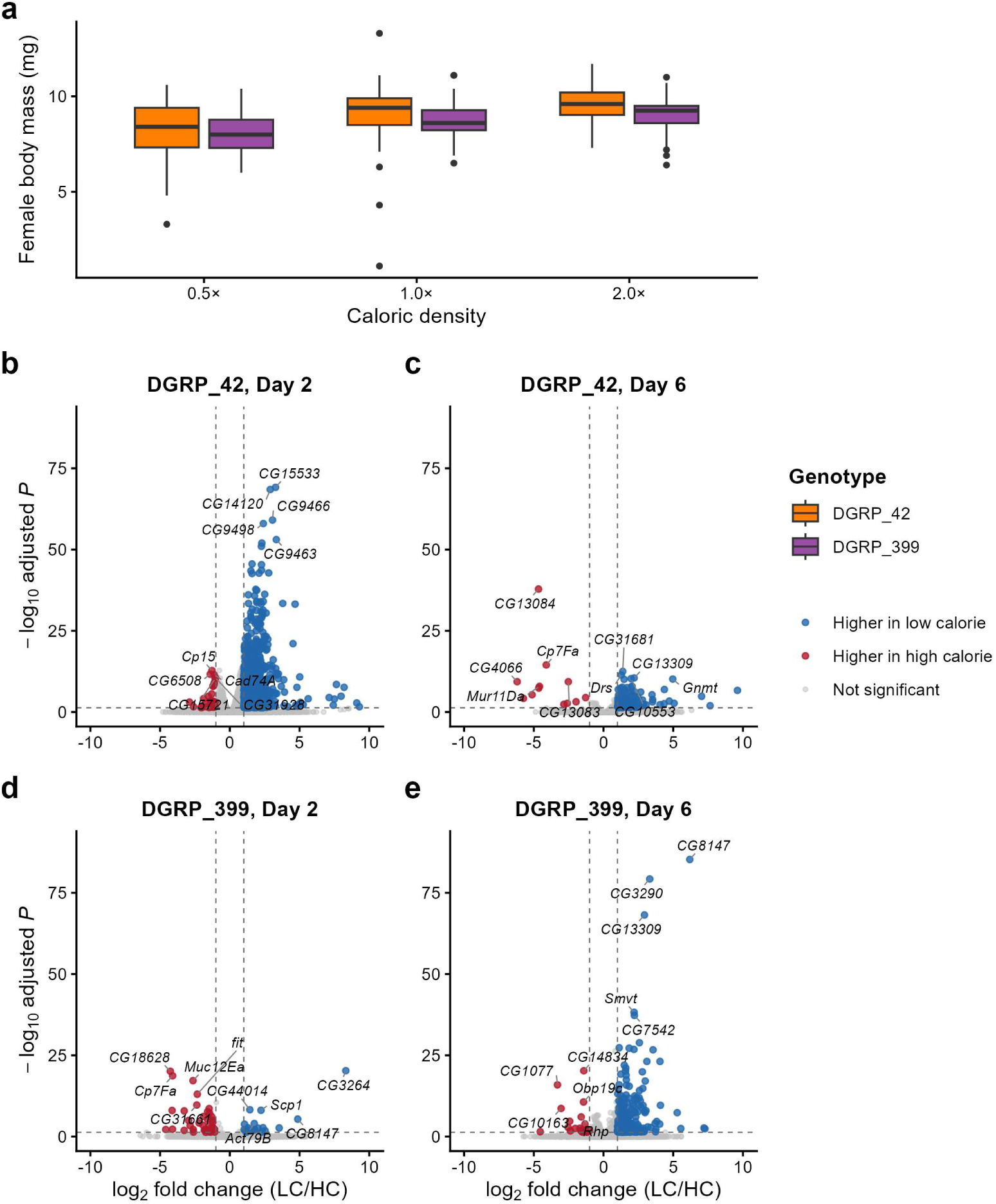
Body mass and transcriptomic responses to caloric density across genetic backgrounds. (a) Adult female body mass (mg) across three dietary treatments (0.5×, 1.0×, and 2.0×) for DGRP_42 and DGRP_399. Horizontal lines indicate medians, boxes represent the interquartile range (IQR), and whiskers extend to the most extreme observations within 1.5 times the IQR. Points beyond the whiskers represent outliers. (b–e) Differential gene expression (DEG) volcano plots. Volcano plots showing statistical significance (− *log*_10_ *adjusted p* -value) versus magnitude of change (*log*_2_ Fold Change) for the extreme dietary contrast (0.5× low-calorie vs. 2.0× high-calorie stress) across genotypes and time points: (b) DGRP_42 Day 2: Robust early transcriptional response to extreme dietary differences. (c) DGRP_42 Day 6: Marked chronological contraction in differentially expressed genes following prolonged exposure. (d) DGRP_399 Day 2: Subtle baseline transcriptional response to dietary divergence. (e) DGRP_399 Day 6: Profound late-stage transcriptomic response, contrasting with the timeline observed in DGRP_42.

**Table 2.** Effects of dietary treatment, genotype, and brood on adult female body mass. Type III ANOVA of a linear mixed-effects model including dietary treatment, genotype, brood, and their two-and three-way interactions as fixed effects, with vial ID as a random effect. Numerator (NumDF) and denominator (DenDF) degrees of freedom, F-statistics, and p-values are reported for each fixed effect. Denominator degrees of freedom were estimated using Satterthwaite’s approximation.

| Term | NumDF | DenDF | F value | *p-value* |
|---|---|---|---|---|
| **Treatment** | **2** | **119.85** | **34.46** | **<0.001** |
| **Genotype** | **1** | **120.05** | **9.17** | **0.003** |
| **Brood** | **6** | **119.25** | **5.72** | **<0.001** |
| Treatment:Genotype | 2 | 119.85 | 0.70 | 0.50 |
| Treatment:Brood | 12 | 118.94 | 1.69 | 0.08 |
| Genotype:Brood | 6 | 119.25 | 1.76 | 0.11 |
| Treatment:Genotype:Brood | 12 | 118.94 | 0.99 | 0.46 |

Tukey-adjusted pairwise comparisons of estimated marginal means revealed that females maintained on the 0.5× diet had significantly lower body mass than those maintained on the 1.0× and 2.0× diets in both genotypes. In DGRP_399, mean female body mass was 8.02, 8.68, and 9.10 mg on the 0.5×, 1.0×, and 2.0× diets, respectively. Females on the 0.5× diet weighed 7.64% less than those on the 1.0× diet (p-value = 0.005) and 11.92% less than those on the 2.0× diet (p-value = 2.67 × 10⁻⁶). Similarly, in DGRP_42, mean female body mass was 8.19, 9.19, and 9.51 mg on the 0.5×, 1.0×, and 2.0× diets, respectively. Females on the 0.5× diet weighed 10.91% less than those on the 1.0× diet (p-value = 1.23 × 10⁻⁵) and 13.92% less than those on the 2.0× diet (p-value = 1.22 × 10⁻⁸). Body mass did not differ significantly between the 1.0× and 2.0× diets in either genotype (DGRP_399: p-value = 0.100; DGRP_42: p-value = 0.268; Supplementary Tables S3 and S4).

### Ovarian Transcripts Confirm Genotype and Diet Specific Responses

Ovarian transcriptomic analysis revealed substantial differences in gene expression between genotypes, dietary treatments, and between timepoints of oogenesis (Table 3). Volcano plots for all pairwise dietary comparisons on Days 2 and 6 are presented in Supplementary Figures S2 and S3 for DGRP_42 and DGRP_399, respectively. The magnitude and timing of the transcriptional response differed markedly between DGRP_42 and DGRP_399, indicating genotype-specific responses to changes in caloric density.

**Table 3.** Summary of differentially expressed genes (DEGs) across diets and timepoints for genotypes DGRP_42 and DGRP_399. This table lists the total number of significantly upregulated and downregulated genes with Benjamini–Hochberg-adjusted p-values < 0.05 and absolute log₂ fold changes > 1 on Day 2 and Day 6 post-exposure. Pairwise contrasts compare High-Calorie (HC; 2.0×) vs. Control (1.0×), Low-Calorie (LC; 0.5×) vs. Control (1.0×), and Low-Calorie vs. High-Calorie (LC vs. HC) conditions. Upregulated and downregulated indicate higher and lower expression, respectively, in the first-listed condition relative to the second.

| comparison |  |  | Upregulated | Downregulated | Total Significant |
|---|---|---|---|---|---|
| 42 | Day2 | HC vs Control | 16 | 105 | 121 |
|  |  | LC vs Control | 310 | 11 | 321 |
|  |  | LC vs HC | 1048 | 29 | 1077 |
|  | Day6 | HC vs Control | 17 | 262 | 279 |
|  |  | LC vs Control | 9 | 3 | 12 |
|  |  | LC vs HC | 143 | 12 | 155 |
| 399 | Day2 | HC vs Control | 14 | 115 | 129 |
|  |  | LC vs Control | 5 | 65 | 70 |
|  |  | LC vs HC | 20 | 39 | 59 |
|  | Day6 | HC vs Control | 27 | 35 | 62 |
|  |  | LC vs Control | 110 | 3 | 113 |
|  |  | LC vs HC | 310 | 24 | 334 |

### DGRP_42 Exhibits an Early Transcriptional Response to Low-Calorie Diet

On Day 2, DGRP_42 exhibited its strongest transcriptional response under low-calorie conditions. The low-calorie (LC) vs. Control comparison identified 321 differentially expressed genes (DEGs), including 310 upregulated and 11 downregulated genes, whereas the LC vs. high-calorie (HC) comparison identified 1,077 DEGs (1,048 upregulated and 29 downregulated), representing the largest transcriptional shift observed in this genotype. In contrast, the HC vs. Control comparison produced only 121 DEGs (16 upregulated and 105 downregulated) (Table 3, Supplementary Figure S4).

Of the 1,077 DEGs identified in the LC vs. HC comparison, 697 were unique to that comparison, whereas 285 genes were shared exclusively between the LC vs. Control and LC vs. HC comparisons, indicating a shared transcriptional response associated with caloric restriction. Only 87 genes were shared exclusively between HC vs. Control and LC vs. HC, while eight genes were common to all three dietary comparisons (Figure 4, Supplementary Figure S4).

**Figure 4.**
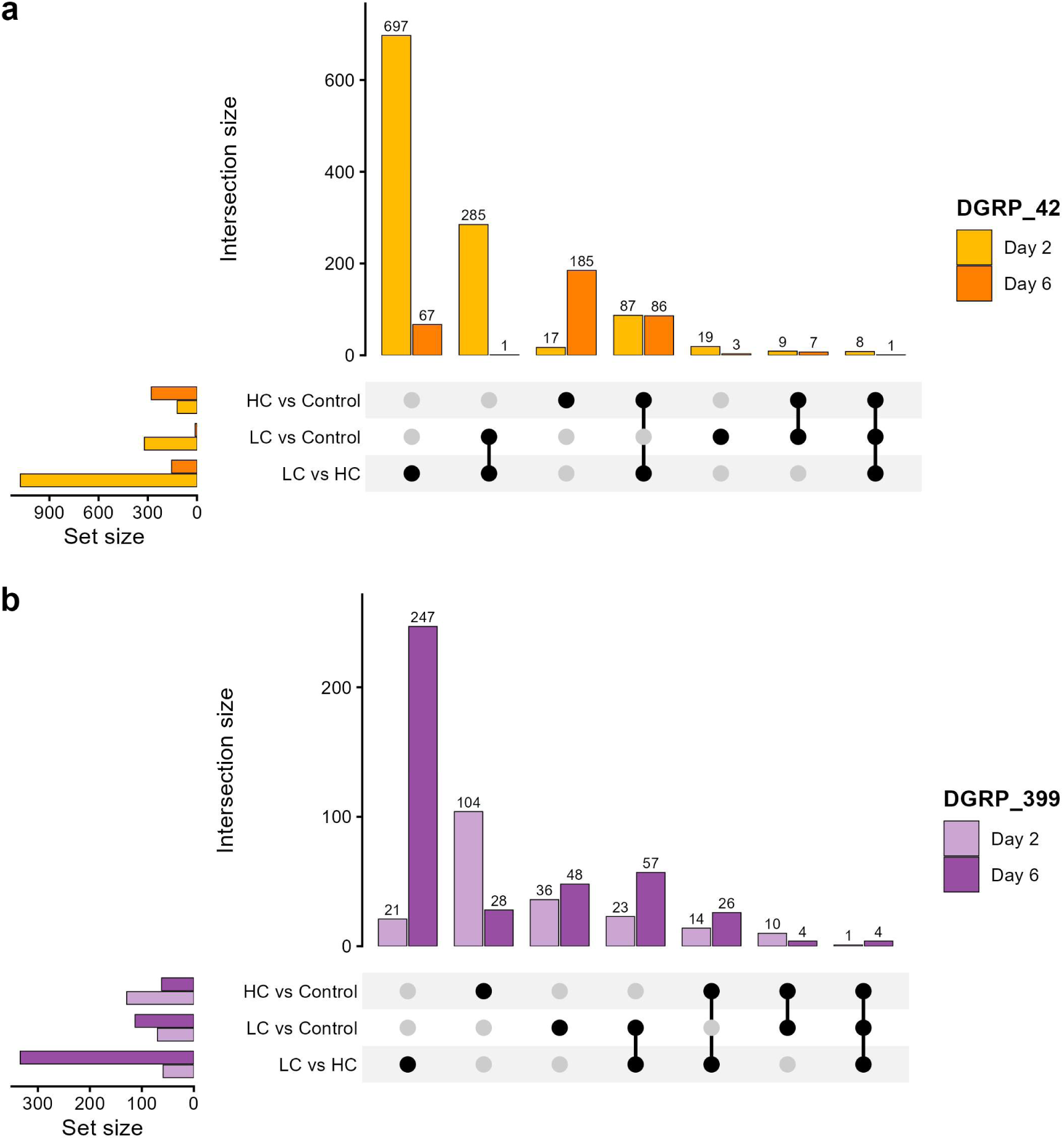
Shared and unique transcriptomic responses to dietary manipulation across timepoints. UpSet plots illustrating the degree of overlap among differentially expressed gene (DEG) sets across dietary contrasts—High Calorie vs. Control (HC_Control), Low Calorie vs. Control (LC_Control), and Low Calorie vs. High Calorie (LC_HC)—evaluated on Day 2 (light shade) and Day 6 (dark shade) for (a) DGRP_42 and (b) DGRP_399. Horizontal bars quantify total DEG set sizes per comparison, while vertical bars detail the number of shared or unique genes within specific contrast combinations across sampling days.

Gene Ontology (GO) enrichment analysis revealed that genes upregulated in LC vs. Control and LC vs. HC were primarily associated with metabolic and enzymatic functions, including drug metabolic process, carbohydrate metabolic process, glucosamine-containing compound metabolic process, amino sugar metabolic process, and chitin metabolic process. Significant enrichment was also observed for GO terms related to transmembrane transport (transporter activity, transmembrane transporter activity, and transmembrane transport), oxidoreductase activity (oxidoreductase activity, heme binding, tetrapyrrole binding, iron ion binding, and cofactor binding), and proteolytic processes (proteolysis, endopeptidase activity, peptidase activity, serine-type endopeptidase activity, and serine hydrolase activity), indicating extensive metabolic remodeling during early caloric restriction. In contrast, genes downregulated predominantly in the HC vs. Control comparison were enriched for GO terms associated with multicellular organismal and developmental processes, chromatin organization (chromosome, chromatin, nucleosome, and protein–DNA complex), membrane transport, and carbohydrate metabolism, suggesting suppression of developmental, structural, and metabolic functions following high-calorie exposure.

Consistent with these findings, KEGG pathway analysis demonstrated that genes upregulated in both the LC vs. Control and LC vs. HC comparisons were enriched for pathways involved in xenobiotic detoxification (Metabolism of xenobiotics by cytochrome P450, Drug metabolism– cytochrome P450, and Drug metabolism–other enzymes), carbohydrate metabolism (Starch and sucrose metabolism, Galactose metabolism, Pentose and glucuronate interconversions, Amino sugar and nucleotide sugar metabolism, and Other glycan degradation), vitamin and cofactor metabolism (Thiamine metabolism, Folate biosynthesis, One carbon pool by folate, Biosynthesis of cofactors, Retinol metabolism, and Ascorbate and aldarate metabolism), lipid and amino acid metabolism (Fatty acid metabolism, Biosynthesis of unsaturated fatty acids, Tyrosine metabolism, Arginine biosynthesis, Glycine, serine and threonine metabolism, and Sphingolipid metabolism), as well as cellular recycling and transport (Lysosome, ABC transporters, and Glutathione metabolism). Additional enrichment was observed for Insect hormone biosynthesis, Neuroactive ligand–receptor interaction, and the Virion–Hepatitis viruses pathway, reflecting broader transcriptional responses to caloric restriction during the early treatment period. In contrast, the HC vs. Control comparison showed comparatively little pathway enrichment, with the Toll and Imd signaling pathway representing the primary significantly upregulated pathway (Figure 5, Supplementary Figure S5).

**Figure 5.**
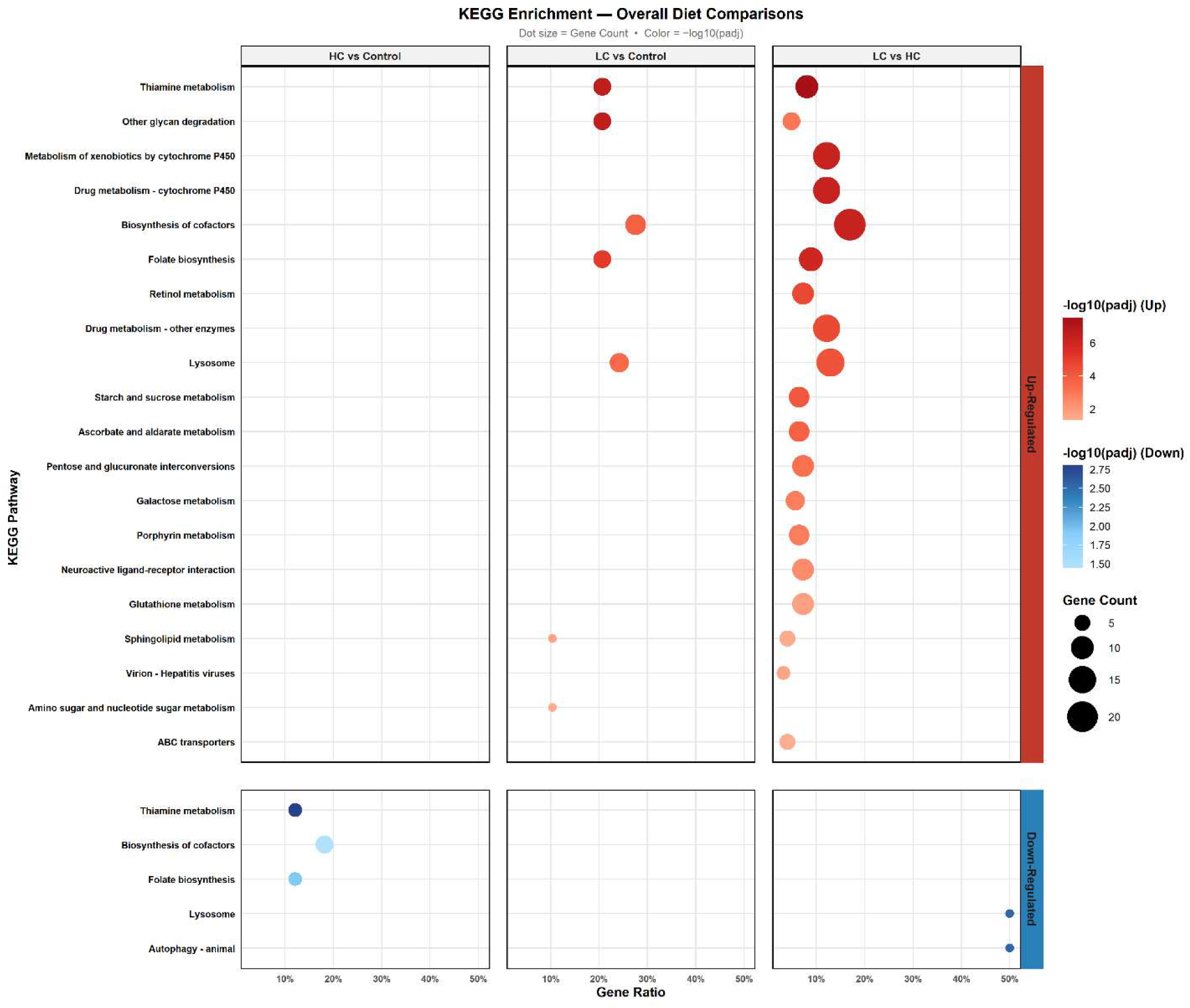
Overall KEGG pathway enrichment analysis across dietary treatments. Bubble plots display significantly enriched KEGG pathways for pairwise dietary comparisons (HC-High caloric vs. Control, LC-low caloric vs. Control, and LC vs. HC) collapsed across genotypes. Direction of regulation is color-coded, with red indicating pathways enriched among upregulated genes and blue indicating pathways enriched among downregulated genes. The x-axis indicates the Gene Ratio, dot size is proportional to the enriched gene count, and color shading reflects statistical significance (darker hues represent higher significance).

By Day 6, the magnitude of transcriptional change shifted markedly toward HC stress. While the response to low-calorie conditions contracted dramatically to only 12 DEGs in the LC vs. Control comparison, the HC vs. Control comparison expanded to 279 DEGs (17 upregulated and 262 downregulated) (Table 3, Supplementary Figure S4).

There were 185 genes unique to HC vs. Control and 86 genes shared exclusively between HC vs. Control and LC vs. HC. The LC vs. Control comparison contained only three unique genes, with seven genes shared with HC vs. Control, one shared with LC vs. HC, and one gene common to all three dietary comparisons (Figure 4, Supplementary Figure S4).

Gene Ontology (GO) enrichment analysis revealed that genes upregulated under low-calorie conditions were predominantly enriched in the LC vs. HC comparison, with overrepresentation of GO terms associated with transmembrane transport (transmembrane transport, transporter activity, transmembrane transporter activity, active transmembrane transporter activity, and primary active transmembrane transporter activity), oxidoreductase activity (oxidoreductase activity, oxidoreductase activity acting on paired donors, heme binding, tetrapyrrole binding, iron ion binding, and cofactor binding), and proteolytic functions (endopeptidase activity, peptidase activity, serine-type peptidase activity, and serine-type endopeptidase activity). Additional enrichment was observed for hydrolase activity (phosphatase activity, phosphoric ester hydrolase activity, and hydrolase activity acting on ester bonds), as well as carbohydrate binding and α-mannosidase activity. In contrast, the LC vs. Control comparison showed limited enrichment, primarily involving ATP-dependent transport processes (ATPase activity, coupled to transmembrane movement of substances, ATPase activity, coupled to movement of substances, and P–P-bond-hydrolysis-driven transmembrane transporter activity), while the HC vs. Control comparison was enriched for developmental GO terms, including multicellular organismal process, multicellular organism development, developmental process, and anatomical structure development. Conversely, genes downregulated under high-calorie conditions were enriched for GO terms related to oxidoreductase activity, transmembrane transport, and proteolytic activity, whereas genes downregulated in the LC vs. HC comparison were primarily associated with chromatin organization and DNA structure, including DNA conformation change, DNA binding, DNA packaging, protein–DNA complex assembly, nucleosome organization, and chromatin assembly or disassembly. No significant GO enrichment was observed among downregulated genes in the LC vs. Control comparison.

KEGG pathway analysis showed that downregulated genes were predominantly enriched for metabolic pathways, including xenobiotic and drug metabolism (Metabolism of xenobiotics by cytochrome P450, Drug metabolism–cytochrome P450, and Drug metabolism–other enzymes), carbohydrate metabolism (Starch and sucrose metabolism, Galactose metabolism, and Pentose and glucuronate interconversions), vitamin and cofactor metabolism (Thiamine metabolism, Folate biosynthesis, Biosynthesis of cofactors, Retinol metabolism, and Ascorbate and aldarate metabolism), cellular recycling and transport (Lysosome, and ABC transporters), amino acid metabolism (Arginine biosynthesis), as well as Neuroactive ligand–receptor interaction. Together, these results indicate broad suppression of metabolic, transport, and nutrient-sensing pathways following prolonged exposure to a high-calorie diet. (Figure 5, Supplementary Figure S5).

### DGRP_399 Exhibits a More Extensive Transcriptional Response to Caloric Restriction on Day 6

On Day 2, DGRP_399 exhibited fewer differentially expressed genes in response to caloric restriction than DGRP_42. In DGRP_399, the HC vs. Control comparison identified 129 DEGs (14 upregulated and 115 downregulated), whereas the LC vs. Control comparison identified 70 DEGs. The LC vs. HC comparison also identified fewer DEGs than in DGRP_42.

Of the 129 DEGs identified in the HC vs. Control comparison, 104 were unique to that comparison, indicating little overlap with genes responding to caloric restriction. The LC vs. Control comparison contained 36 unique genes, while 23 genes were shared exclusively between LC vs. Control and LC vs. HC. Only one gene was common to all three dietary comparisons (Figure 4, Supplementary Figure S4).

Gene Ontology (GO) enrichment analysis indicated a comparatively modest transcriptional response in DGRP_399 on Day 2. Genes upregulated under low-calorie conditions were primarily enriched in the LC vs. HC comparison for biological processes associated with carbohydrate and carbohydrate-derivative metabolism (carbohydrate derivative metabolic process, glucosamine-containing compound metabolic process, aminoglycan metabolic process, amino sugar metabolic process, and chitin metabolic process), lipid metabolism (lipid metabolic process and transferase activity, transferring acyl groups), and hydrolase activity (hydrolase activity, acting on ester bonds), with additional enrichment for chitin binding. In the LC vs. Control comparison, enrichment was limited to phosphoric ester hydrolase activity, phosphatase activity, structural molecule activity, and structural constituent of cuticle, whereas no significant GO enrichment was detected in the HC vs. Control comparison. In contrast, genes downregulated under HC vs. Control were enriched for GO terms related to ATP-dependent and transmembrane transport (ATPase activity, ATPase activity coupled to transmembrane movement of substances, P–P-bond-hydrolysis-driven transmembrane transporter activity, transporter activity, and transmembrane transport), oxidoreductase activity (oxidoreductase activity, oxidoreductase activity acting on paired donors, heme binding, tetrapyrrole binding, iron ion binding, and cofactor binding), and transition metal ion binding. Downregulated genes in the LC vs. Control comparison were enriched primarily for proteolytic processes, including proteolysis, endopeptidase activity, peptidase activity, serine-type endopeptidase activity, serine hydrolase activity, metallopeptidase activity, and metalloendopeptidase activity, while the LC vs. HC comparison showed enrichment for only a single GO term, odorant binding.

In Day 6 ovaries, DGRP_399 appeared to mount a stronger response to the low-calorie diet than on Day 2. The LC vs. Control comparison identified 113 DEGs (110 upregulated and 3 downregulated), whereas the LC vs. HC comparison expanded to 334 DEGs (310 upregulated and 24 downregulated) (Table 3, Supplementary Figure S4).

There was extensive overlap between low-calorie comparisons, with 57 genes shared exclusively between LC vs. Control and LC vs. HC, while the LC vs. HC comparison contained 247 unique genes. In contrast, the HC vs. Control comparison contracted to only 28 unique genes, and four genes were shared across all three dietary comparisons (Figure 4, Supplementary Figure S4)

Gene Ontology (GO) enrichment analysis revealed upregulated genes under low-calorie conditions, particularly in the LC vs. HC comparison and largely shared with the LC vs. Control comparison, were enriched for biological processes associated with carbohydrate and lipid metabolism (carbohydrate metabolic process, monosaccharide metabolic process, hexose metabolic process, and lipid metabolic process), glycosyl transfer and hydrolysis (transferase activity, transferring glycosyl groups, transferase activity, transferring hexosyl groups, hydrolase activity acting on glycosyl bonds, and hydrolase activity hydrolyzing O-glycosyl compounds), oxidoreductase activity (oxidoreductase activity, oxidoreductase activity acting on paired donors, heme binding, tetrapyrrole binding, iron ion binding, and cofactor binding), proteolytic processes (proteolysis, peptidase activity, endopeptidase activity, serine-type peptidase activity, serine-type endopeptidase activity, and metallopeptidase activity), and transmembrane transport (transporter activity, transmembrane transporter activity, and transmembrane transport). Additional enrichment was observed for chitin binding, carbohydrate binding, mannosidase activity, α-mannosidase activity, phosphatase activity, and phosphoric ester hydrolase activity. Although a subset of these GO terms was also significantly enriched in the HC vs. Control comparison, they were represented by fewer genes and lower levels of significance. In contrast, genes downregulated under HC vs. Control were predominantly enriched for GO terms related to drug and carbohydrate metabolism (drug metabolic process, carbohydrate metabolic process, glucosamine-containing compound metabolic process, amino sugar metabolic process, aminoglycan metabolic process, and chitin metabolic process), transmembrane transport, extracellular region, proteolytic activity, and chitin binding. The LC vs. HC comparison showed additional enrichment for transferase activity, transferring sulfur-containing groups and sulfotransferase activity, whereas no unique major functional categories were observed beyond those already enriched in the HC vs. Control comparison.

KEGG pathway analysis showed that genes upregulated under low-calorie conditions (primarily LC vs. Control and LC vs. HC) were enriched for pathways involved in vitamin and cofactor metabolism (Thiamine metabolism, Biosynthesis of cofactors, Folate biosynthesis), xenobiotic detoxification (Drug metabolism–cytochrome P450, Metabolism of xenobiotics by cytochrome P450, and Drug metabolism–other enzymes), carbohydrate metabolism (Starch and sucrose metabolism, Galactose metabolism, Pentose and glucuronate interconversions, and Other glycan degradation), lipid and amino acid metabolism (Glycerophospholipid metabolism, Sphingolipid metabolism, and Arginine biosynthesis), as well as lysosomal function, glutathione metabolism, retinol metabolism, porphyrin metabolism, Toll and Imd signaling, and Neuroactive ligand– receptor interaction. In contrast, the HC vs. Control comparison exhibited minimal pathway enrichment, with only Oxidative phosphorylation, Autophagy – animal, and Lysosome significantly upregulated only in day 6 and not in day 2 (Supplementary Figure S6).

On Day 2, genes downregulated in the HC vs. Control comparison were enriched for pathways associated with vitamin and cofactor metabolism (Thiamine metabolism, Biosynthesis of cofactors, and Folate biosynthesis), carbohydrate metabolism (Starch and sucrose metabolism), and ABC transporters. In the LC vs. Control comparison, downregulated genes were enriched for Lysosome, Hormone signaling, and Glycosaminoglycan degradation, whereas the LC vs. HC comparison showed enrichment for Lysosome and Autophagy – animal. By Day 6, genes downregulated in the HC vs. Control comparison were enriched for Galactose metabolism, Pentose and glucuronate interconversions, One carbon pool by folate, Glycine, serine and threonine metabolism, and the Toll and Imd signaling pathway (Supplementary Figure S6).

Enrichment of Autophagy – animal among downregulated genes was also observed in the overall LC vs. HC comparison (Figure 5).

### Conserved Transcriptional Responses Across Genotypes

Despite their contrasting temporal responses between Day 2 and Day 6, both DGRP_42 and DGRP_399 shared a conserved transcriptional signature for a subset of genes. Across both exposure periods, *CG4783* and *Smvt* (*Sodium-dependent multivitamin transporter*) were consistently upregulated in LC conditions and downregulated in HC conditions.

Overall, these results demonstrate distinct temporal signatures of transcriptional regulation between the two genotypes to the same caloric density gradient. DGRP_42 mounted a rapid response to caloric restriction that shifted toward high-calorie stress over time, whereas DGRP_399 exhibited the opposite pattern, responding initially to excess calories before mounting a delayed but robust transcriptional response to caloric restriction

### Caloric Restriction Reduces Fecundity in Marker-Present but Not Marker-Free Crosses

In marker-free crosses, fecundity was not significantly affected by dietary treatment (*F*₂,₁₈ = 1.53, *p*-value = 0.243), genotype (*F*₁,₁₈ = 1.36, *p*-value = 0.260), or their interaction (*F*₂,₁₈ = 1.28, *p*-value = 0.301; Table 4). Tukey-adjusted pairwise comparisons within each genotype detected no significant differences in fecundity among dietary treatments (all *p*-values ≥ 0.102; Figure 6a; Supplementary Tables S5 and S6).

**Table 4.** Effects of dietary treatment, genotype, and brood on female fecundity. Type III F-tests with Satterthwaite-approximated denominator degrees of freedom were obtained from linear mixed-effects models fitted separately to marker-free and marker-present crosses. Treatment, genotype, brood, and their interactions were included as fixed effects, with vial identity as a random intercept. Fecundity was measured as eggs per female per 24 h. DF represents numerator degrees of freedom, and Sum Sq represents Type III sums of squares. Denominator degrees of freedom ranged from 18 to 108 for marker-free crosses and from 88 to 528 for marker-present crosses. Significant effects (p-value < 0.05) are indicated in bold.

| Term | DF | **Marker free** |  | **Marker present** |  |
|---|---|---|---|---|---|
|  |  | Sum Sq | *p*-value | Sum Sq | *p*-value |
| Treatment | 2 | 619.60 | 0.24 | **610.41** | **<0.001** |
| Genotype | 1 | 273.93 | 0.26 | **1901.60** | **<0.001** |
| Brood | 6 | 363.18 | 0.94 | **8392.58** | **<0.001** |
| Treatment: Genotype | 2 | 518.49 | 0.30 | 141.03 | 0.12 |
| Treatment: Brood | 12 | 3014.62 | 0.26 | **1025.98** | **0.002** |
| Genotype: Brood | 6 | 1049.75 | 0.52 | **985.58** | **<0.001** |
| Treatment: Genotype: Brood | 12 | 1401.40 | 0.86 | 320.72 | 0.62 |

**Figure 6.**
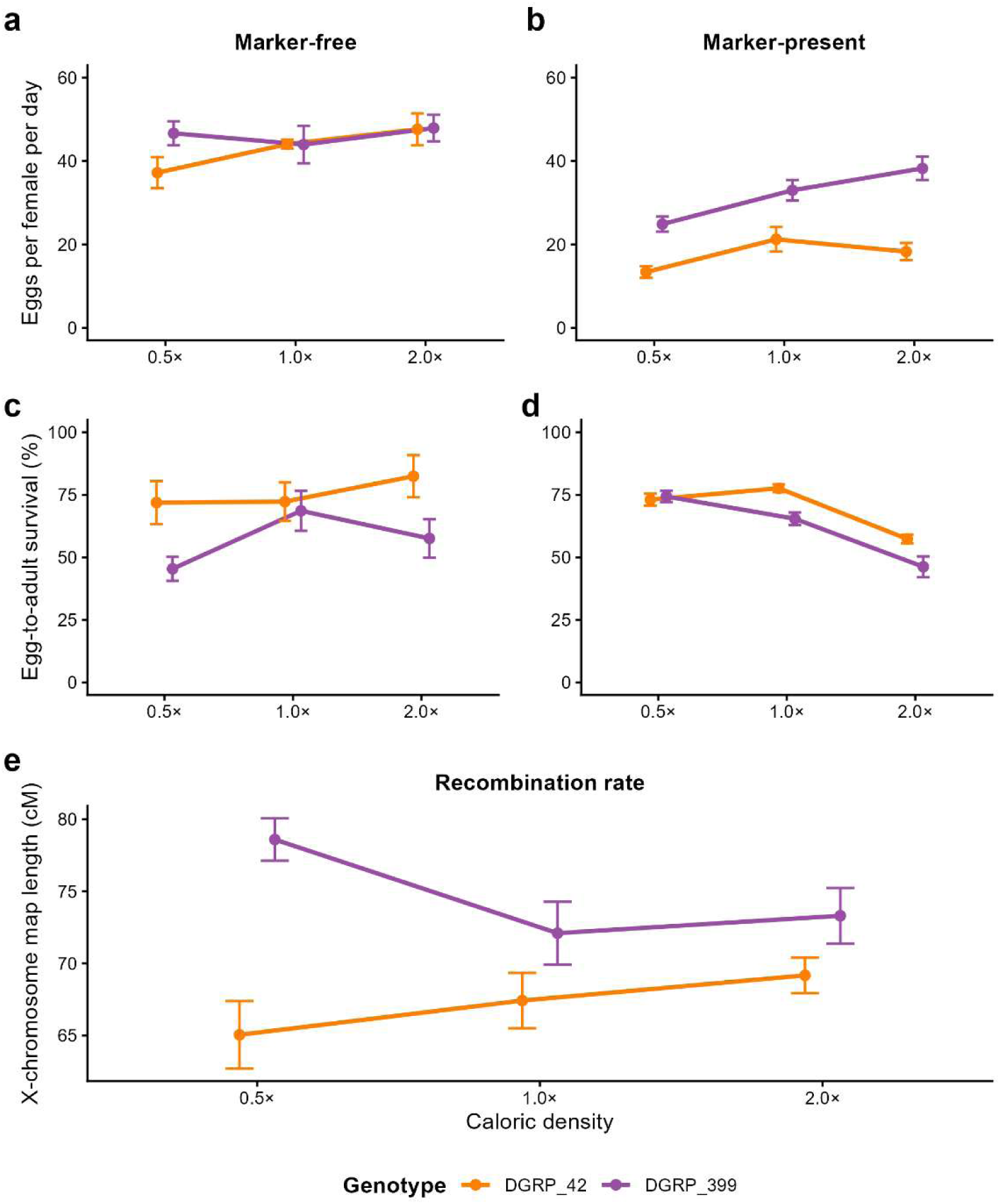
X-chromosome meiotic recombination rates and fitness phenotypes across a nutritional gradient. (a) Mean daily maternal fecundity (eggs per female per 24 h) in marker-free crosses across three dietary treatments (0.5×, 1.0×, and 2.0×) for DGRP_42 and DGRP_399. (b) Mean daily maternal fecundity (eggs per female per 24 h) in marker-present crosses across the same dietary treatments and genotypes. (c) Marker-free egg-to-adult viability (%) isolating natural offspring survival responses to caloric restriction and excess across genotypes. (d) Marker-present egg-to-adult viability (%) detailing survival patterns influenced by visible marker allele combinations. These viability measurements were used to account for the potential effects of differential offspring survival on estimates of meiotic recombination. (e) Viability-corrected X-chromosome genetic map lengths (cM) across dietary treatments for DGRP_42 and DGRP_399.

In marker-present crosses, fecundity was significantly affected by dietary treatment (*F*₂,₈₈ = 9.45, *p*-value = 0.000192) and genotype (*F*₁,₈₈ = 58.85, *p*-value = 2.16 × 10⁻¹¹), with DGRP_399 producing more eggs per female per day (24.9–38.2 across diets) than DGRP_42 (13.4–21.2 across diets; Figure 6b). The treatment-by-genotype interaction was not significant (*F*₂,₈₈ = 2.18, *p*-value = 0.119). Fecundity also varied significantly among broods (*F*₆,₅₂₈ = 43.29, *p*-value < 0.0001), with significant genotype-by-brood (*F*₆,₅₂₈ = 5.08, *p*-value < 0.0001) and treatment-by-brood interactions (*F*₁₂,₅₂₈ = 2.65, *p*-value = 0.00189; Table 4).

In marker-present crosses, fecundity was significantly lower on the 0.5× diet than on the 1.0× control diet in both genotypes. In DGRP_42, fecundity decreased by 37.0% (*p*-value = 0.0439), whereas in DGRP_399, fecundity decreased by 24.6% (*p*-value = 0.0397). Fecundity was also 35.0% lower on the 0.5× diet than on the 2.0× diet in DGRP_399 (*p*-value = 0.000281), but this comparison was not significant in DGRP_42 (*p*-value = 0.286). Neither genotype exhibited a significant difference in fecundity between the 1.0× and 2.0× diets (DGRP_42: *p*-value = 0.632; DGRP_399: *p*-value = 0.258; Supplementary Tables S5 and S6).

### Dietary Effects on Egg-to-Adult Survival were Detected in Marker-Present but Not Marker-Free Crosses

In marker-free crosses, egg-to-adult survival was significantly affected by genotype (*F*₁,₁₇ = 8.80, *p*-value = 0.0087) and brood (*F*₆,₁₀₈ = 4.69, *p*-value < 0.001), whereas neither dietary treatment (*F*₂,₁₇ = 1.56, *p*-value = 0.239) nor the treatment-by-genotype interaction (*F*₂,₁₇ = 1.41, *p*-value = 0.271) was significant (Figure 6, Table 5). In DGRP_399, estimated marginal mean survival was 45.4%, 68.6%, and 57.6% on the 0.5×, 1.0×, and 2.0× diets, respectively. Relative to the 1.0× control diet, survival was numerically lower by 33.8% on the 0.5× diet and 16.1% on the 2.0× diet. In DGRP_42, estimated marginal mean survival was 71.9%, 72.3%, and 82.4% on the 0.5×, 1.0×, and 2.0× diets, respectively, corresponding to a 0.6% decrease on the 0.5× diet and a 14.0% increase on the 2.0× diet relative to the control. None of the pairwise dietary contrasts within either genotype reached statistical significance after Tukey adjustment (all *p*-values ≥ 0.106; Supplementary Tables S7 and S8).

**Table 5.** Egg-to-adult survival in the marker-free and marker-present cohorts Type III analysis of variance for linear mixed-effects models (LMMs).Results of generalized linear mixed models testing the effects of Treatment (diet), Genotype, Brood, and their interactions on egg-to-adult survival, fit separately for the marker-free and marker-present datasets. Significance was assessed using Type III tests with Satterthwaite’s method for denominator degrees of freedom (df); vial ID and data collector were included as random effects. Sum Sq = Type III sum of squares; DF = numerator degrees of freedom for each term; bold/significant terms are those with Pr(>F) < 0.05.

| Term | DF | **Marker free** |  | **Marker present** |  |
|---|---|---|---|---|---|
|  |  | Sum Sq | *p*-value | Sum Sq | *p*-value |
| Treatment | 2 | 1373.68 | 0.24 | **16718.32** | **<0.001** |
| Genotype | 1 | **3874.78** | **0.01** | **2139.90** | **0.001** |
| Brood | 6 | **12399.21** | **<0.001** | **9809.51** | **<0.001** |
| Treatment: Genotype | 2 | 1244.66 | 0.27 | **1591.47** | **0.012** |
| Treatment: Brood | 12 | 3312.14 | 0.82 | 2302.69 | 0.34 |
| Genotype: Brood | 6 | 1133.94 | 0.86 | 1109.34 | 0.37 |
| Treatment: Genotype: Brood | 12 | 7887.29 | 0.14 | 2377.46 | 0.31 |

In marker-present crosses, egg-to-adult survival was significantly affected by dietary treatment (*F*₂,₈₂.₂ = 48.82, *p*-value = 1.06 × 10⁻¹⁴), genotype (*F*₁,₈₂.₂ = 12.50, *p*-value = 0.00067), and brood (*F*₆,₅₂₂.₀ = 9.55, *p*-value = 5.66 × 10⁻¹⁰), with a significant treatment-by-genotype interaction (*F*₂,₈₂.₂ = 4.65, *p*-value = 0.0122; Table 5). In DGRP_399, estimated marginal mean survival declined with increasing caloric density, from 74.4% on the 0.5× diet to 65.4% on the 1.0× diet and 47.9% on the 2.0× diet. Survival on the 0.5× diet was 13.6% higher than on the 1.0× diet (*p*-value = 0.0248), whereas survival on the 2.0× diet was 26.8% lower than on the 1.0× diet (*p*-value = 4.23 × 10⁻⁶) and 35.5% lower than on the 0.5× diet (*p*-value = 3.06 × 10⁻¹⁰). In DGRP_42, estimated marginal mean survival was 73.1%, 77.6%, and 57.4% on the 0.5×, 1.0×, and 2.0× diets, respectively. Survival did not differ significantly between the 0.5× and 1.0× diets (*p*-value = 0.348), but survival on the 2.0× diet was 26.1% lower than on the 1.0× diet (*p*-value = 6.65 × 10⁻⁸) and 21.5% lower than on the 0.5× diet (*p*-value = 2.18 × 10⁻⁵; Supplementary Tables S7 and S8).

Both genotypes exhibited comparable reductions in egg-to-adult survival on the high-calorie diet relative to the control (26.8% in DGRP_399 and 26.1% in DGRP_42). However, the two genotypes exhibited contrasting responses to caloric restriction: survival increased significantly in DGRP_399, whereas no significant change was detected in DGRP_42. These patterns are consistent with the significant treatment-by-genotype interaction observed in marker-present crosses.

### Effects of Genotype, Diet, and Brood on Genetic Map Length

Mean recombination rates by genotype and treatment are summarized in Table 1. Overall, DGRP_399 exhibited longer genetic map lengths than DGRP_42 across all diet treatments (Table 1, **Figure 6**e). This rank order was consistent with (Hunter, Huang, et al. 2016), who reported higher mean *yellow–vermilion* recombination frequencies in DGRP_399 (32.05%) than in DGRP_42 (30.80%), although their estimates covered a shorter X-chromosome interval.

Mean Kosambi map lengths for DGRP_42 were 69.36, 69.54, and 72.13 cM for the 0.5×, 1.0×, and 2.0× diets, respectively. Corresponding values for DGRP_399 were 79.78, 72.92, and 77.09 cM. Analysis of Kosambi map lengths detected significant effects of genotype (F₁,₈₃.₁ = 12.93, *p-value =* 0.00055) and brood (F₆,₅₀₉.₁ = 3.28, *p-value =* 0.0035), together with a significant Genotype × Brood interaction (F₆,₅₀₉.₂ = 3.00, *p-value =* 0.0068). Neither the main effect of dietary treatment (F₂,₈₃.₁ = 1.56, *p-value =* 0.217) nor the Genotype × Treatment interaction (F₂,₈₃.₂ = 1.50, *p-value =* 0.228) was significant. The Brood × Treatment interaction (F₁₂,₅₀₉.₁ = 1.01, *p-value =* 0.437) and the three-way Genotype × Treatment × Brood interaction (F₁₂,₅₀₉.₁ = 1.42, *p-value =* 0.154) were also nonsignificant (Supplementary Table S9).

Mean viability-corrected genetic map lengths estimated using the extended counting model were 65.86, 67.55, and 69.20 cM for DGRP_42 on the 0.5×, 1.0×, and 2.0× diets, respectively. Corresponding estimates for DGRP_399 were 78.61, 72.10, and 73.26 cM (Table 1). Analysis of viability-corrected map lengths detected a significant effect of genotype (F₁,₈₃.₇ = 23.81, *p-value =* 5.01 × 10⁻⁶), together with significant Genotype × Treatment (F₂,₈₃.₉ = 3.73, *p-value =* 0.028) and Genotype × Brood interactions (F₆,₅₁₂.₆ = 4.20, *p-value =* 0.00039). Neither dietary treatment (F₂,₈₃.₈ = 0.80, *p-value =* 0.452) nor brood (F₆,₅₁₂.₆ = 1.21, *p-value =* 0.300) had a significant main effect. The three-way Genotype × Treatment × Brood interaction was not significant (F₁₂,₅₁₂.₆ = 0.96, *p-value =* 0.484; Table 7). Brood-specific variation in viability-corrected genetic map length across genotypes and dietary treatments is illustrated in Supplementary Figure S7.

Pairwise comparisons of estimated marginal means revealed that, in DGRP_399, the viability-corrected genetic map length was 8.8% greater on the 0.5× diet (78.54 cM) than on the 1.0× diet (72.19 cM; *p*-value = 0.038). Genetic map length did not differ significantly between the 0.5×and 2.0× diets (*p*-value = 0.099) or between the 1.0× and 2.0× diets (*p*-value = 0.915). No pairwise dietary contrasts within DGRP_42 were significant (all *p*-values > 0.36; Supplementary Tables S10 and S11).

Interval-specific recombination rates for each genotype and dietary treatment combination are presented in Table 6. The *f–v* interval consistently exhibited the highest recombination rates across genotypes and dietary treatments, whereas the *y⁺–car* and *car–f* intervals exhibited the lowest rates. The *v–cv* and *cv–sc* intervals exhibited intermediate recombination rates. Despite variation in total genetic map length among genotypes and dietary treatments, the relative distribution of recombination across the assayed X-chromosome intervals was broadly similar among experimental groups. Interval-specific differences between maximum-likelihood estimates and observed recombination fractions are presented in Supplementary Figure S8.

**Table 6.** Interval-specific meiotic recombination fractions along the Drosophila melanogaster X-chromosome across dietary treatments and genotypes. Mean viability-corrected recombination fraction estimates (± SE) are presented for five adjacent intervals defined by visible genetic markers (y⁺–car, car–f, f–v, v–cv, and cv–sc) spanning 98% of the X chromosome. Estimates are summarized across three dietary treatments (0.5× low-calorie, 1.0× control, and 2.0× high-calorie) for DGRP_42 and DGRP_399.

|  | **DGRP_42** |  |  | **DGRP_399** |  |  |
|---|---|---|---|---|---|---|
| interval | 0.5x | 1.0x | 2.0x | 0.5x | 1.0x | 2.0x |
| *y+-car* | 0.059 ± 0.005 | 0.055 ± 0.005 | 0.076 ± 0.007 | 0.071 ± 0.004 | 0.076 ± 0.005 | 0.069 ± 0.005 |
| *car-f* | 0.074 ± 0.005 | 0.072 ± 0.005 | 0.093 ± 0.007 | 0.089 ± 0.004 | 0.087 ± 0.004 | 0.082 ± 0.004 |
| *f-v* | 0.242 ± 0.011 | 0.251 ± 0.010 | 0.242 ± 0.011 | 0.266 ± 0.007 | 0.266 ± 0.007 | 0.254 ± 0.007 |
| *v-cv* | 0.161 ± 0.009 | 0.156 ± 0.007 | 0.164 ± 0.010 | 0.221 ± 0.007 | 0.187 ± 0.008 | 0.202 ± 0.009 |
| *cv-sc* | 0.124 ± 0.007 | 0.140 ± 0.007 | 0.117 ± 0.008 | 0.139 ± 0.006 | 0.104 ± 0.006 | 0.125 ± 0.006 |

**Table 7.** Viability-corrected X-chromosome meiotic recombination rate generalized linear mixed model (GLMM) evaluating the effects of diet, genotype, and brood. The model evaluates the effects of dietary treatment, genotype, and brood, as well as their two- and three-way interactions, on viability-corrected recombination rate estimated using the Koury et al. (2026) framework. The viability correction accounts for differential survival among progeny classes, allowing recombination to be evaluated while accounting for post-meiotic viability differences that may vary among genotypes and dietary environments. Numerator and denominator degrees of freedom (NumDF and DenDF), F-statistics, and associated p-values are reported for each fixed effect. Significant effects are indicated in bold (p-value < 0.05). The model detected significant effects of genotype, Treatment × Genotype, and Genotype × Brood. For the corresponding analysis of Kosambi map lengths, see Supplementary Table S9.

| Term | DF | DenDF | Sum Sq | Mean Sq | F value | *p*-value |
|---|---|---|---|---|---|---|
| Treatment | 2 | 83.76 | 306.90 | 153.45 | 0.80 | 0.45 |
| **Genotype** | **1** | **83.75** | **4,560.24** | **4,560.24** | **23.81** | **<0.001** |
| Brood | 6 | 512.63 | 1,389.71 | 231.62 | 1.21 | 0.30 |
| **Treatment: Genotype** | **2** | **83.87** | **1,426.99** | **713.50** | **3.73** | **0.03** |
| Treatment: Brood | 12 | 512.58 | 3,310.93 | 275.91 | 1.44 | 0.14 |
| **Genotype: Brood** | **6** | **512.61** | **4,827.78** | **804.63** | **4.20** | **<0.001** |
| Treatment: Genotype: Brood | 12 | 512.59 | 2,212.76 | 184.40 | 0.96 | 0.48 |

### No Significant Association Between Recombination Rate and Egg-to-Adult Survival

Dietary responses of egg-to-adult survival and viability-corrected genetic map length did not consistently correspond to one another in marker-present crosses. In DGRP_399, both egg-to-adult survival and genetic map length were significantly greater on the 0.5× diet than on the 1.0× diet. However, although both genotypes exhibited significantly reduced survival on the 2.0× diet relative to the control, neither exhibited a significant difference in genetic map length between these treatments. In DGRP_42, genetic map length did not differ significantly among dietary treatments despite significant dietary differences in survival (Supplementary Tables S7, S8, S10, and S11).

To further examine the relationship between recombination rate and offspring survival, we fitted separate linear mixed-effects models for each genotype, dietary treatment, and brood combination, with data collector included as a random effect. No statistically significant association between egg-to-adult survival and recombination rate was detected across the experiment (all *p*-values ≥ 0.0796; Figure 7). The highest positive slope was observed in DGRP_42 on the 0.5× diet during brood A (β = 0.633, SE = 0.358, *p*-value = 0.099), whereas the highest negative slope was observed in DGRP_399 on the 0.5× diet during brood D (β = −0.508, SE = 0.322, *p*-value = 0.139).

**Figure 7.**
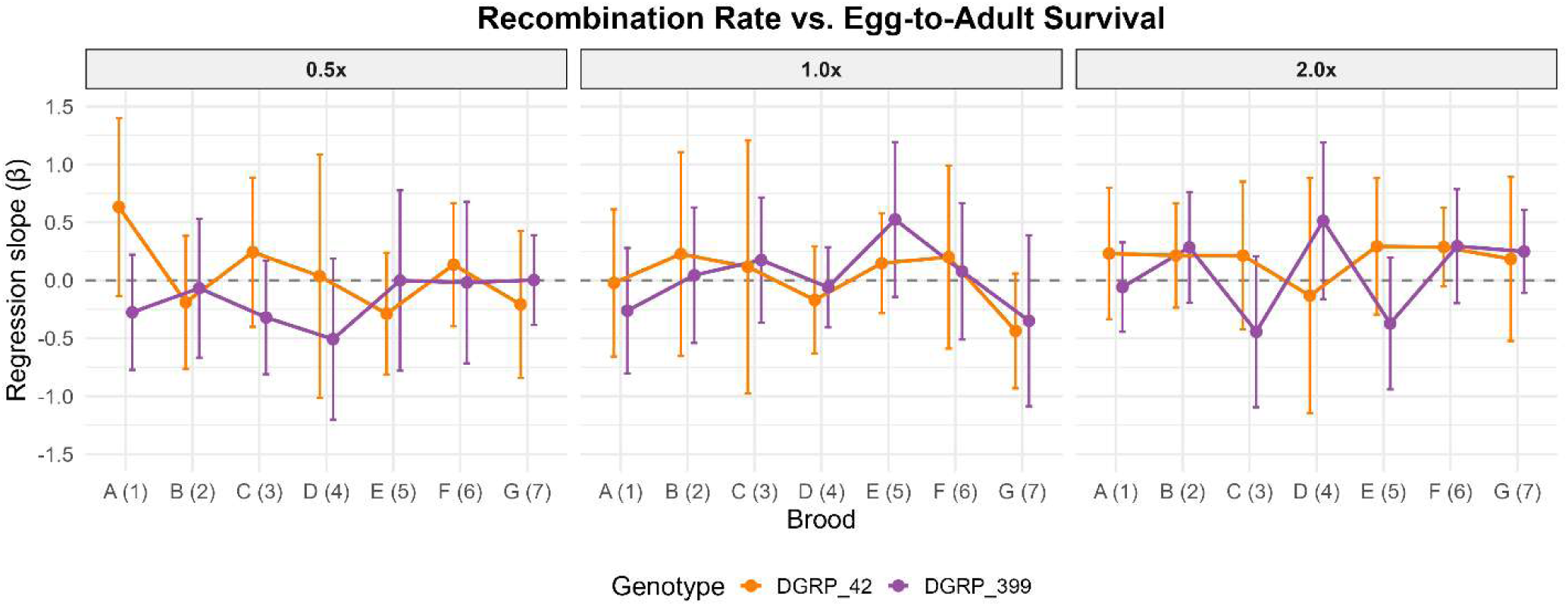
Association between meiotic recombination rate and egg-to-adult survival across dietary treatments. Estimated regression slopes (β) from linear mixed-effects models fitted separately for each genotype (DGRP_42 and DGRP_399), dietary treatment (0.5×, 1.0×, and 2.0×), and brood in marker-present crosses, with data collector included as a random effect. Points represent estimated regression slopes, and error bars indicate 95% confidence intervals. The horizontal dashed line indicates a slope of zero. Broods A–G correspond to seven consecutive 24-hour egg-laying periods (Days 1–7).

Overall, despite significant treatment-by-genotype interactions for both egg-to-adult survival and viability-corrected genetic map length, we did not detect significant associations between recombination rate and offspring survival within genotype, dietary treatment, and brood combinations

### No Significant Effects of Diet or Genotype on Crossover Interference

Mean interference ranged from 2.75 (DGRP_399, 1.0× diet) to 4.75 (DGRP_399, 2.0× diet), with DGRP_42 showing more stable estimates across diets (3.25–3.5) than genotype DGRP_399 (2.75–4.75) (Table 1). Neither diet (F₂,₁₈ = 1.23, *p-value =* 0.315) nor genotype (F₁,₁₈ = 0.17, *p-value =* 0.689) had a significant effect on interference, and the Diet × Genotype interaction was also not significant (F₂,₁₈ = 0.72, *p-value =* 0.501). Residual variance was substantial relative to the between-group sums of squares (residual MS = 2.26 vs. total explained MS = 1.6–2.8 across terms), consistent with the modest sample size (*n* = 4 replicate estimates per genotype × dietary treatment combination) may have limited statistical power to detect differences in crossover interference.

## Discussion

Dietary manipulation across a gradient of caloric density (0.5×–2.0×) affected adult female body mass and ovarian transcriptional profiles in *Drosophila melanogaster*, contributing to genotype-dependent responses in meiotic recombination rates. Across both genetic backgrounds (DGRP_42 and DGRP_399), mean female body mass increased with dietary caloric density, with a significant main effect of dietary treatment (*F*₂,₁₁₉.₈₅ = 34.46, *p*-value = 1.50 × 10⁻¹²). While both genotypes exhibited similar body mass responses to dietary variation, their ovarian transcriptional responses differed temporally. DGRP_42 exhibited a more extensive transcriptional response to caloric restriction on Day 2, whereas DGRP_399 exhibited a more extensive response on Day 6 than on Day 2. Furthermore, viability-corrected genetic map lengths exhibited a significant genotype-by-treatment interaction despite the absence of a significant main effect of dietary treatment.

By incorporating experimental mortality and marker-associated differential viability directly into recombination rate estimates (Koury et al. 2026), our results suggest that nutritional environments affect recombination rates in genotype-dependent ways. Historically, classic Drosophila testcrosses evaluated crossover rates solely from surviving F_2_ adults. Differential survival among offspring classes can bias the proportions of haplotypes observed among surviving progeny, potentially affecting estimates of meiotic recombination (Koury et al. 2026). Without controlling for experimental mortality, viability distortions can mimic or obscure true changes in crossover frequency (Kong et al. 2004; Koury et al. 2026). When accounting for post-meiotic viability, we detected a significant Genotype-by-Environment (G × E) interaction in X-chromosome meiotic recombination rates. Notably, traditional Kosambi-corrected estimates failed to reach statistical significance for this interaction (*p*-value >0.05), suggesting that accounting for offspring viability can improve the power to detect genotype-dependent recombination responses.

Both analytical approaches detected a significant genotype effect and a Genotype × Brood interaction. However, the Genotype × Diet interaction was significant only in the analysis of viability-corrected map lengths, and the main effect of brood was significant only in the Kosambi analysis. These differences highlight the importance of accounting for progeny survival when interpreting recombination responses to dietary variation. Because differential survival among offspring classes can alter the marker combinations observed among surviving adults, observed recombination fractions may reflect both meiotic recombination and post-meiotic survival. We therefore used viability-corrected estimates to evaluate associations between recombination and reproductive performance. This correction accounts for modeled differences in progeny viability but does not itself establish a causal relationship between recombination and fitness.

Alongside genotype-dependent recombination responses to diet, we evaluated associations between recombination rate and offspring survival within 42 genotype-by-diet-by-brood combinations, comprising two genetic backgrounds, three dietary treatments, and seven brood periods. Effect sizes were small, confidence intervals broadly overlapped zero, and no association reached statistical significance. Thus, we found no consistent support for the negative association between fitness and recombination predicted under fitness-associated recombination (FAR) in the conditions examined. These results suggest that the recombination rate plasticity observed here is not adaptive, but we cannot exclude an intermediate association with aspects of physiological condition not captured by offspring survival or measured in our experiment.

Hadany and Beker (2003) proposed that an association between fitness and recombination could be mediated by factors such as stress responses, age, or DNA damage. Consequently, the absence of an association with offspring survival does not exclude a relationship between recombination and an intermediate physiological correlate of fitness. Our study examined only two DGRP backgrounds derived from a single natural population (Mackay et al. 2012), which may not encompass the genetic variation needed to detect a broader fitness–recombination relationship.

Furthermore, the dietary treatments used here were selected to maintain sufficient offspring for recombination estimation and may have imposed insufficient physiological stress to reveal a negative association. Together, these limitations restrict our conclusions to the genetic backgrounds, dietary conditions, and fitness components examined. In contrast to the changes in recombination rate, analysis of crossover interference revealed no significant differences across diets or genotypes, nor a significant G × E interaction. This suggests that dietary effects on recombination frequency were not accompanied by detectable changes in crossover interference under the conditions examined (Zhao et al. 1995; Koury et al. 2026).

By incorporating experimental mortality into recombination estimates, our study provides evidence that dietary effects on meiotic recombination depend on genetic background, suggesting these responses do not reflect adaptive adjustments to reduced fitness.

### Diet Induces Shared Metabolic Responses with Genotype-Specific Timing

The temporal differences in ovarian transcriptional responses between DGRP_42 and DGRP_399 may be relevant to the brood-specific variation in meiotic recombination rates, although a direct relationship between these responses was not tested. Ovarian transcriptomes were sampled on Days 2 and 6 from virgin F1 females separate from those used for recombination assays. Although both groups experienced the same dietary treatments, mating status represents an important difference between the two experimental groups. Mating can alter ovarian physiology, including germline stem-cell proliferation and egg production, through neuroendocrine signaling (Ameku and Niwa 2016). Furthermore, because RNA-seq was performed on whole ovaries, the observed transcriptional differences may reflect changes in the relative abundance of ovarian cell types and developmental stages, as well as changes in gene expression within individual cells. Therefore, the Day 2 and Day 6 transcriptomes characterize ovarian responses to prolonged dietary exposure in virgin females but may not fully represent the ovarian molecular environment of mated females used in recombination assays. Previous work demonstrates that nutritional conditions dynamically regulate the Drosophila ovary, affecting germline and somatic stem-cell proliferation, egg-chamber development, and cell survival, with insulin signaling contributing substantially to these responses (Drummond-Barbosa and Spradling 2001; Drummond-Barbosa 2008). Thus, the temporal differences observed between DGRP_42 and DGRP_399 are consistent with the broader capacity of the Drosophila ovary to dynamically adjust its developmental and physiological state in response to nutritional conditions.

DGRP_42 mounted a robust early transcriptional response to caloric restriction on Day 2, but this response was largely reduced by Day 6, indicating a less extensive transcriptional response by Day 6. In contrast, DGRP_399 exhibited a delayed transcriptional response, particularly under caloric restriction, indicating a more prolonged molecular adjustment. This delayed response coincided with elevated recombination frequencies in later broods of DGRP_399, which may have originated from oocytes developing within an ovarian environment represented more closely by the Day 6 transcriptome (Supplementary Figure S7).

Although the transcriptomic and recombination datasets cannot be directly linked to individual oocytes, the temporal correspondence between delayed transcriptional changes and altered recombination is consistent with the possibility that sustained nutritional signaling influences the developmental environment in which meiotic recombination is established. Nutritional regulation of the Drosophila germline is known to occur through multiple physiological and endocrine pathways, including insulin-dependent mechanisms that alter germline stem-cell proliferation and cyst development responses (Drummond-Barbosa and Spradling 2001; Drummond-Barbosa 2008). Conversely, despite the extensive early transcriptional response in DGRP_42, recombination frequencies remained comparatively stable across broods, suggesting that this genotype may either buffer the effects of dietary stress more effectively or that early transcriptional changes were insufficient to alter crossover outcomes or we have not captured due to the recombination assay design. Together, these findings indicate that the timing of the ovarian molecular response to dietary stress differs between genotypes and may contribute to their contrasting patterns of recombination plasticity.

The enrichment of metabolic and nutrient-responsive pathways provides additional support for a physiological connection between dietary state and ovarian function. In DGRP_42, the early transcriptional response was enriched for metabolic and detoxification pathways, including cytochrome P450 and glutathione S-transferase pathways, together with components of insulin/insulin-like signaling (IIS) and Target of Rapamycin (TOR) signaling. In contrast, DGRP_399 exhibited a delayed metabolic response, particularly under caloric restriction. IIS and TOR are established nutrient-sensing pathways in Drosophila and are important regulators of germline stem-cell activity, ovarian growth, and oogenesis (Drummond-Barbosa and Spradling 2001; Lin and Hsu 2020). In particular, TOR signaling has been linked directly to nutrient-mediated ovarian growth through the TOR–Spargel/dPGC-1 pathway, demonstrating that nutritional information can be transmitted into the ovary to regulate egg-chamber development and female fertility (Abul Basar et al. 2019).

Specific gene responses further connect these transcriptional patterns to dietary metabolism. Glycine N-methyltransferase (*Gnmt*) expression was higher in LC than HC conditions in both genotypes on both days, with log₂ fold changes ranging from 2.53 to 4.97 and all comparisons meeting our DEG criteria. *Gnmt* participates in S-adenosylmethionine metabolism, and previous experiments demonstrated that its function contributes to the lifespan response to dietary restriction in Drosophila (Obata and Miura 2015). Its consistent response here suggests that methionine-related metabolism is a shared component of the ovarian response to dietary variation, although transcript abundance alone does not establish changes in metabolic flux or methylation.

Carbohydrate-related regulation provides another connection to previous dietary studies. In DGRP_42 on Day 2, target of brain insulin (*tobi*) expression was higher in LC than HC conditions (log₂FC = 1.86, adjusted *p-value =* 1.75 × 10⁻³⁸). Expression was also lower in HC than control conditions, whereas LC and control did not differ significantly. Buch et al. (2008)demonstrated that dietary protein and sugar regulate *tobi* in opposite directions through insulin- and glucagon-like signaling. This finding highlights the importance of dietary composition alongside total nutrient availability. However, because our treatments varied food concentration rather than independently manipulating protein and carbohydrate, the observed response cannot be attributed to a particular nutrient. The LC–HC difference in *tobi* expression was not statistically significant in the other genotype–time combinations.

Literature-guided examination of nutrient-response genes also identified smaller expression changes that met the adjusted *p*-value threshold but fell below our twofold-change criterion. Thor, which encodes the translational regulator 4E-BP, showed lower expression in HC than control conditions on Day 2 in both DGRP_42 (log₂FC = −0.47, adjusted *p-value =* 0.0044) and DGRP_399 (log₂FC = −0.81, adjusted *p-value =* 0.0098). In DGRP_42, Thor expression was also higher in LC than HC conditions (log₂FC = 0.82, adjusted *p-value =* 2.09 × 10⁻⁸). These directions are consistent with previous evidence linking dietary restriction to 4E-BP regulation and its roles in translational control and mitochondrial function (Zid et al. 2009; Kang et al. 2017). Nevertheless, these modest transcript changes do not establish altered 4E-BP protein activity or downstream translation.

Similarly, *GstD1* expression was higher in LC than HC conditions in DGRP_42 on Day 6 (log₂FC = 0.62, adjusted *p-value =* 0.037). (Brookheart et al. 2017) reported reduced ovarian antioxidant-gene expression, including *GstD1*, following high-sucrose feeding. Our result is directionally consistent with that observation, although increasing sucrose and changing overall dietary concentration are distinct nutritional manipulations. This comparison supports considering redox regulation as a possible component of the ovarian dietary response, rather than demonstrating oxidative damage.

Despite their contrasting temporal responses, both genotypes also shared higher expression of CG4783 and Sodium-dependent multivitamin transporter (*Smvt*) in LC than HC conditions on both days. Both genes met our DEG criteria in all four genotype–time comparisons, although differences from the control diet were not uniformly significant. Experimental work in Drosophila has established a role for Smvt-mediated biotin uptake in intestinal stem-cell function (Neophytou and Pitsouli 2022), providing a functional basis for investigating whether its ovarian response involves micronutrient transport. Our results do not, however, demonstrate increased vitamin uptake or establish a nutrient-sensing mechanism. CG4783 remains functionally uncharacterized, and its consistent dietary response identifies it as a candidate for further investigation.

We also examined genes included in the supplementary analysis of Novak et al. (2026) of piRNA-associated and chromatin regulators. In DGRP_42 on Day 2, *AGO3* and *mael* expression was lower in LC than HC conditions (log₂FC = −0.37 for each; adjusted *p-value =* 0.027 and 0.0043, respectively). Although neither gene met our twofold-change criterion for differential expression, these results suggest that dietary conditions may influence the expression of genes associated with piRNA-mediated regulation.

These pathways therefore provide a plausible physiological route through which dietary conditions could alter the developmental environment of the germline. However, the present data do not establish a direct causal pathway from IIS or TOR signaling to meiotic double-strand break formation or crossover designation. Meiotic recombination in Drosophila is initiated through programmed DNA double-strand breaks involving *MEI-P22* and *MEI-W68*, while subsequent crossover formation is regulated by meiosis-specific factors including the mei-MCM complex (Liu et al. 2002; Hartmann et al. 2019). Therefore, the enrichment of nutrient-responsive pathways in our ovarian transcriptomes should be interpreted as evidence that dietary stress alters the molecular environment in which germline development and meiosis occur, rather than as direct evidence that these pathways control crossover formation. This distinction is important because the observed genotype-specific temporal responses may reflect differences in how each genetic background senses, buffers, or transduces nutritional signals during oogenesis.

The enrichment of detoxification and antioxidant pathways may provide a second potential connection between nutritional state and germline function. Dietary perturbations have been shown to alter ovarian metabolism, antioxidant gene expression, mitochondrial function, and oxidative damage in Drosophila Brookheart et al. (2017). Moreover, experimentally induced oxidative stress in Drosophila oocytes can cause premature loss of cohesion and increased chromosome-segregation errors during meiosis, demonstrating that the redox environment can affect meiotic chromosome integrity (Perkins et al. 2016). Thus, the genotype-specific enrichment of detoxification and glutathione-associated pathways observed here may indicate differences in the ability of the two backgrounds to buffer nutritional perturbations at the cellular level. Whether such changes directly influence crossover formation or instead affect later meiotic chromosome stability remains an important question for future work.

These findings extend previous evidence that dietary manipulation produces genotype-dependent ovarian transcriptional responses. Novak et al. (2026) reported diet-associated enrichment of amino-acid metabolism, carbohydrate metabolism, and oxidation–reduction processes, with a more extensive transcriptional response in DGRP_42 than DGRP_217. Our gene-level findings similarly implicate nutritional metabolism, while revealing responses shared between genetic backgrounds alongside differences associated with genotype and sampling time. Together, these findings motivate experiments that distinguish the effects of dietary concentration from nutrient composition, which can independently influence *Drosophila* reproduction and longevity (Lee et al. 2008; Grandison et al. 2009). Whether the transcriptional responses identified here contribute to recombination plasticity or represent parallel physiological effects of diet remains unresolved.

### No Consistent Relationship Between Recombination Rate Changes and Reproductive Fitness

Fitness-associated recombination (FAR) describes a negative association between an individual’s fitness and its recombination rate, potentially preserving fitter genetic combinations while increasing the shuffling of less-fit combinations (Hadany and Beker 2003; Agrawal et al. 2005). We evaluated the prediction that lower reproductive performance is associated with higher recombination rates. Specifically, we tested whether recombination rate was negatively associated with maternal fecundity or offspring survival across the dietary conditions examined.

Our results did not reveal a consistent negative association between recombination rate and reproductive fitness. As shown in **Figure 7**, regression slopes across time periods did not consistently fall below zero; instead, their directions and magnitudes varied among genotypes, treatments, and brood periods. In addition to offspring survival, maternal fecundity provides a biologically relevant proxy for reproductive performance. Similar to survival, our fecundity measurements also did not exhibit a consistent negative relationship with recombination rate. For example, although both genotypes exhibited significantly reduced fecundity under caloric restriction in marker-present crosses, DGRP_42 and DGRP_399 displayed contrasting recombination reaction norms across the same dietary treatments. These findings provide no consistent support for the tested FAR prediction across the dietary conditions examined, although they do not rule out fitness-associated recombination under other conditions.

Several features of the experimental design limit the strength of this inference. Fecundity and offspring survival represent components of reproductive fitness rather than complete measures of maternal fitness (Orr 2009). Furthermore, fitness components and recombination were measured at the vial level, with three F_1_ females per vial, so the analysis evaluates associations among experimental units rather than within individual females. The dietary treatments were selected to maintain sufficient offspring for reliable recombination estimates, which may have limited the range of fitness reduction examined. Consequently, the absence of statistically significant associations should not be interpreted as evidence that biologically meaningful relationships are absent, particularly where estimates are imprecise.

Importantly, our results do not imply that fitness-dependent recombination is impossible. Previous work in Drosophila provides evidence that recombination responses can depend on the physiological sensitivity of the genotype to environmental stress. Under desiccation stress, desiccation-sensitive genotypes exhibited stronger changes in recombination rate and crossover interference than desiccation-tolerant genotypes, with the authors interpreting these responses as evidence for fitness-dependent recombination plasticity (Aggarwal et al. 2019). Differences between those findings and our nutritional results may reflect variation in stressor type, genetic background, treatment severity, or experimental design.

Although the relationship between recombination and the measured fitness components remains uncertain, our results demonstrate that recombination responses to dietary variation differ between the genetic backgrounds examined. This interpretation is consistent with previous work demonstrating that genetic background significantly contributes to variation in recombination rate in Drosophila and that interactions between genotype and physiological state can alter the magnitude and direction of recombination responses (Hunter, Huang, et al. 2016). More broadly, recent theoretical work emphasizes that recombination plasticity can arise from complex interactions among environmental stress, genetic background, and individual fitness (Rybnikov et al. 2023).

### Differences in Experimental Design May Explain Contrasting Dietary Responses

To evaluate the impact of caloric density on meiotic recombination, we first compared our findings alongside (Novak et al. 2026) which previously examined dietary effects using the same DGRP_42 genotype. Although both studies used DGRP_42, differences in thermal environment, genetic architecture, the chromosomal regions examined, and the methods used to account for offspring viability may partly explain the differences in our findings.

First, Novak et al. (2026) conducted their assays at 25°C, whereas our experiment was maintained at 21°C. This 4°C difference is important because temperature is a well-established environmental determinant of meiotic recombination in *Drosophila melanogaster* (Plough 1917; Grell 1978; Stevison et al. 2017). Classic work demonstrated that temperature can alter crossover frequency, while subsequent studies have shown that environmental temperature can generate plastic responses in recombination rate and crossover patterning (Smith 1936; Stevison et al. 2017). Therefore, differences in ambient temperature between the two studies could contribute to differences in the magnitude or direction of the observed dietary response.

Second, the genetic composition of tested flies differed significantly between the two studies. In (Novak et al. 2026), testcross F_1_ females inherited a 50% genome-wide DGRP_42 background across all chromosomes. In contrast, our experimental crossing design reduced the expected DGRP genetic contribution to 25% across both the X chromosome and autosomes, while recombination was specifically assessed along the X chromosome. By restricting DGRP background contribution, our experimental design minimized background epistatic noise from autosomal variation and isolated chromosome-specific genetic influences on meiotic exchange. Additionally, the crossing design limited inbreeding effects by reducing homozygosity and the associated expression of recessive deleterious alleles that could otherwise contribute to offspring mortality (Koury et al. 2026).

Furthermore, while Novak et al. (2026) measured recombination using four markers spanning the region between the *yellow* and *forked* loci of the X chromosome, our study employed six markers spanning approximately 98% of the X-chromosome genetic map (Koury et al. 2026), providing broader coverage of meiotic crossover patterns along this chromosome.

Finally, the two studies diverged in their analytical objectives. (Novak et al. 2026) focused primarily on organismal physiological changes. In contrast, our study centered on Genotype-by-Environment (G × E) interactions, empirical fitness proxies (maternal fecundity and egg-to-adult survival), and the relationship between fitness loss and recombination rates. Even when raw distance metrics (such as Kosambi mapping) display similar overall directional trends in DGRP_42 across diets, uncorrected recombination fractions fail to separate genuine meiotic plasticity from differential zygotic mortality. By pairing marker-present testcrosses with marker-free control crosses under the extended *C*(*x*)*M* counting model (Koury et al. 2026), we accounted for differential post-meiotic viability. These results are consistent with genotype-dependent recombination reaction norms in response to dietary variation.

### Evolutionary Implications of Genotype-Specific Recombination Plasticity

The significant genotype-by-environment (G × E) interaction observed in meiotic recombination rates has important implications for understanding the evolutionary consequences of environmental variation. Because natural environments fluctuate in resource availability, genotype-specific recombination plasticity may influence how existing genetic variation is reshuffled among offspring. However, the consequences of these responses for adaptation and longer-term evolutionary change remain to be determined. This interpretation is consistent with evidence that genetic background is an important determinant of recombination rate and that interactions between genotype and physiological state can generate heterogeneous recombination reaction norms in *Drosophila* (Hunter, Robinson, et al. 2016). Similarly, dietary manipulation has been shown to influence meiotic crossover frequency in a genotype-dependent manner in male mice (Belmonte-Tebar et al. 2022). Together with our findings, these studies highlight the importance of genetic background in mediating recombination plasticity across diverse environmental and physiological conditions.

Experimental evolution studies in *D. melanogaster* have demonstrated that populations exposed to different thermal regimes can diverge in recombination rate without corresponding divergence in temperature-associated recombination plasticity, suggesting that the genetic architectures underlying these traits may be distinct (Kohl and Singh 2018). Further investigation of these populations revealed divergence in fine-scale recombination landscapes despite broader-scale conservation, suggesting that recombination modifiers may respond to selection during evolutionary change (Winbush and Singh 2022). Together, these findings highlight the importance of distinguishing environmentally induced recombination plasticity from evolutionary changes in recombination rate.

The observed genotype-specific recombination responses do not establish whether this plasticity is adaptive. Such responses could have beneficial, neutral, or detrimental consequences, depending on their effects on offspring genotypes and reproductive success. Recent syntheses of the recombination-plasticity literature indicate that environmental perturbations can alter recombination rates; however, the evolutionary significance of these changes remains unresolved and may depend on both genetic background and the specific characteristics of the environmental challenge (Rybnikov et al. 2023). The contrasting responses of DGRP_42 and DGRP_399 to identical dietary environments shown here further suggest that nutritional changes do not induce a consistent directional change in recombination across genetic backgrounds.

Further experiments are needed to determine whether fitness-associated recombination depends on stressor type, treatment severity, genetic background, or developmental timing. Future experiments incorporating a wider range of dietary conditions, including shifts in protein to carbohydrate ratio, additional genetic backgrounds, and other stressors could help determine when negative fitness–recombination associations occur and how precisely they can be estimated.

## Data availability statement

Raw RNA sequencing (RNA-seq) data generated in this study have been deposited in the National Center for Biotechnology Information (NCBI) Gene Expression Omnibus (GEO) under accession number GSE353957 (https://www.ncbi.nlm.nih.gov/geo/query/acc.cgi?acc=GSE353957).

All other raw data and code for all analyses have been integrated into a GitHub repository with a static DOI (will request upon acceptance) which can be found at https://github.com/StevisonLab/Caloric-Density-and-Recombination.

## Acknowledgments

We would like to thank members of the Stevison Lab for helpful feedback throughout this project. Stocks obtained from the Bloomington Drosophila Stock Center (NIH P40OD018537) were used in this study. This work was completed in part with resources provided by the Auburn University Easley Cluster. The authors acknowledge the use of Large Language Models to assist in the preparation of this manuscript. Specifically, Google’s Gemini 1.5 Pro and NotebookLM were utilized for text editing, grammatical refinement, and summarizing reference articles. Additionally, Anthropic’s Claude 3.5 Sonnet and OpenAI’s GPT-5.3-mini were used for code generation and debugging via functional prompts. The authors maintain full responsibility for the accuracy of the final text, the validity of the generated code, and the absence of plagiarism.

